# Lipopolysaccharide lateral mobility in the Gram-negative bacterial outer membrane is confined and governed by interactions within the conserved Lipid A anchor

**DOI:** 10.1101/2025.03.10.642448

**Authors:** Joe Nabarro, Rosalyn M. Leaman, Samuel Lenton, Leonore Mantion, Richard J. Spears, Mark C. Coles, Dmitri O. Pushkin, Martin A. Fascione, Christoph G. Baumann

## Abstract

The Gram-negative bacterial cell envelope is defined by an asymmetric outer membrane where the outer leaflet adopts a highly ordered structure composed principally of lipopolysaccharide molecules. The organisation and dynamics of these glycolipids are key to the ability of the outer membrane to act as an innate barrier against chemical and antibiotic challenges, and as a load bearing element for the cell. Strong intermolecular forces are thought to govern the lateral diffusion of lipopolysaccharide in the outer membrane, but the extent and molecular basis of this diffusion has remained a controversial topic for over 50 years. Here we use a bio-orthogonal labelling strategy and *in vivo* fluorescence microscopy to unequivocally demonstrate extreme lateral confinement of lipopolysaccharide in the outer membrane of *Escherichia coli*, regardless of carbohydrate domain size and structure. We specifically identify magnesium cation-mediated interactions at the base of the carbohydrate and hydrophobic interactions within the lipid milieu as critical for lipopolysaccharide confinement. Importantly, these traits are conserved across multiple pathogenic species irrespective of O-antigen and capsular serotype. Together, these findings establish lipopolysaccharide endotoxin lateral confinement as a ubiquitous feature of the outer membrane and highlight potential universal vulnerabilities of the bacterial cell envelope.

## Introduction

Outer membrane (OM) asymmetry is critical to cellular integrity and antibiotic resistance in Gram-negative bacteria ^1,2^, with several redundant molecular systems employed to actively maintain it ^3^. The amphipathic glycolipid lipopolysaccharide (LPS) is a primary component of the OM, alongside β-barrel outer membrane proteins (OMPs) ^4,5^, and as such its role in governing OM morphology and function is well established ^6,7^. Lipid A anchors LPS in the membrane and has a chemical structure conserved across most Gram-negative bacteria ^8^, comprising a bis-phosphorylated β-1,6-glucosamine disaccharide bearing six acyl chains (Figure 1A,D and Supplementary Figure 1). The hydrophobic elements result in the tight packing of LPS promoted by intermolecular van der Waals forces that maintain the lipid component in the outer leaflet of the OM in a gel-like state with low fluidity ^9^, in sharp contrast to a typical fluid phospholipid membrane bilayer. Despite its significance in the OM, the lateral mobility of LPS and the molecular interactions underpinning its well-established functional role remain underexplored.

**Figure 1:**
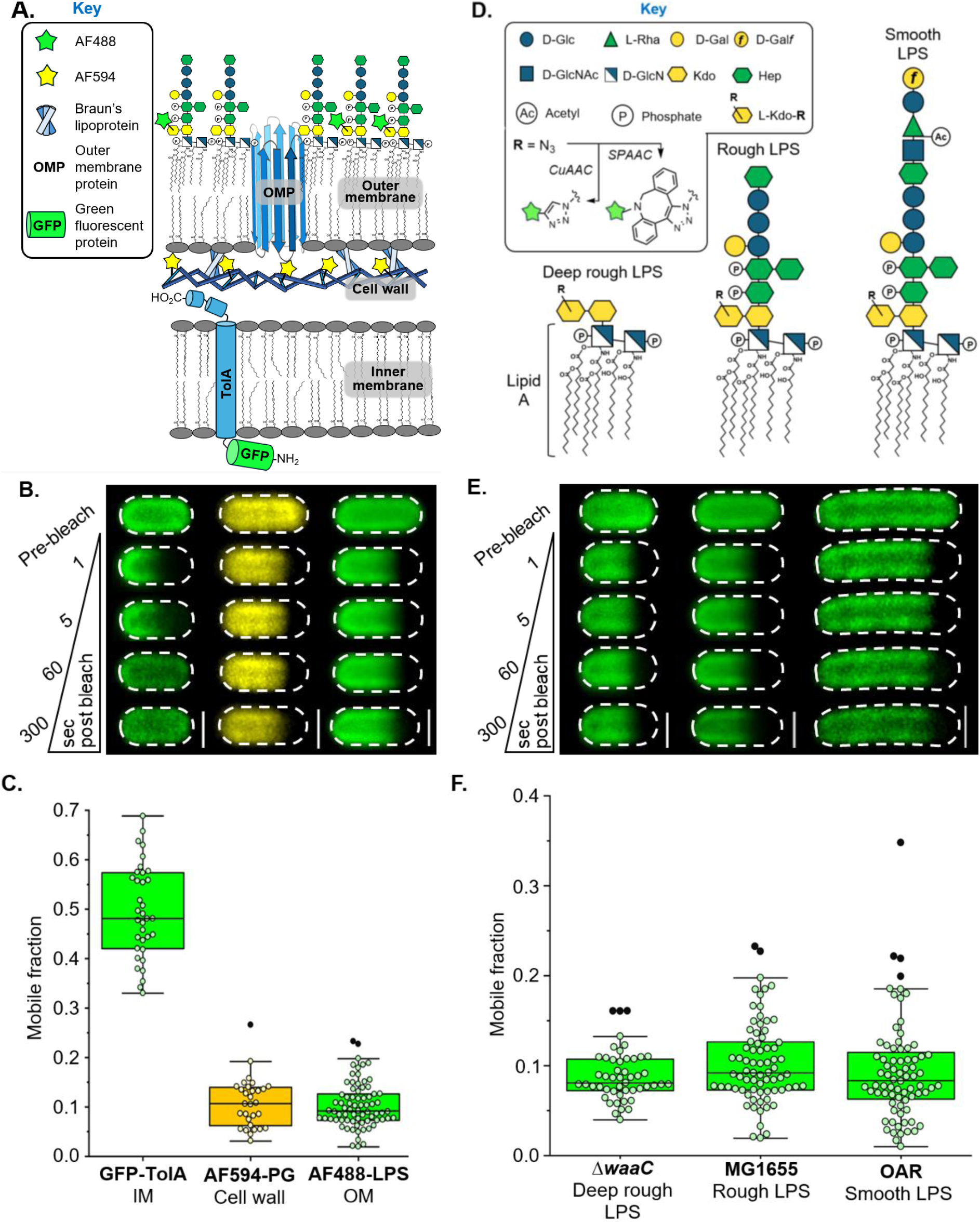
LPS lateral mobility in the OM is very restricted and influenced by intermolecular interactions involving the conserved Lipid A-(Kdo)_2_ core. **A.** Schematic of bacterial cell envelope showing specific targets which were fluorescently labelled with an organic dye or GFP to assess lateral mobility by confocal fluorescence recovery after photobleaching (FRAP) microscopy. Cell wall peptidoglycan (PG) and the transmembrane TolA complex are characterised by very restricted and mostly unrestricted lateral mobilities, respectively. Fluorescent-labelling of LPS via a 2-step bio-orthogonal approach enabled its lateral mobility to be assessed via FRAP. **B.** Representative FRAP sequences of GFP-TolA in the IM (left), AF594-PG in the periplasm (middle), and AF488-labelled LPS in the OM outer leaflet (right) of *E. coli* cells. Scale bars: 1.0 µm. **C.** Comparison of mobile fractions for GFP-TolA (median = 0.476, n = 36), AF594-PG (median = 0.107, n = 30) and AF488-LPS (median = 0.090, n = 69) demonstrate that LPS lateral mobility is very restricted. Each symbol represents an individual measurement with black symbols classified as outliers. **D.** Schematics of deep rough, rough and smooth LPS glycoforms present in Δ*waaC*, MG1655 and O-antigen restored (OAR) *E. coli* strains, respectively, showing the composition of the oligosaccharide domain displayed at the cell surface. *In situ* metabolic labelling of the chemically conserved LPS inner core oligosaccharide domain was done with an azide functionalised Kdo-analogue which enabled specific conjugation of alkyne-functionalised organic fluorescent dyes via Cu(I)-catalysed (CuAAC) or Cu(I)-free strain promoted azide-alkyne cycloaddition (SPAAC). **E.** Representative FRAP sequences for AF488-LPS in the OM of deep rough LPS-producing (left), rough LPS-producing (middle), and smooth LPS-producing (right) *E. coli* cells. Scale bars: 1.0 µm. **F.** Comparison of FRAP-derived mobile fractions show that restriction of LPS lateral mobility is independent of the different glycoforms present in Δ*waaC* (median = 0.081, n = 49), MG1655 (median = 0.091, n = 69) and OAR (median = 0.083, n = 75) *E. coli* strains.

Divalent cation-mediated electrostatic interactions between conserved anionic groups exposed at the base of LPS have long been hypothesised to influence its diffusion and therefore OM organisation ^10,11^. Additionally, intermolecular hydrogen bonds mediated by the more distal regions of the oligosaccharide domain may also play a role ^12^. These intermolecular interactions are predicted to restrict LPS lateral mobility in the OM contributing to the tight packing which underpins its barrier function ^13,14^. However, only a limited number of *in vivo* studies have attempted to characterise the lateral mobility of LPS in the OM of intact cells ^15–17^, with contrasting findings. Mühlradt *et al*. (1974) used pulse-chase experiments and electron microscopy to measure the average mobility of ferritin-labelled smooth LPS in the OM of *Salmonella enterica* serovar Typhimurium, concluding LPS lateral mobility was tightly confined ^15^. This was subsequently contradicted by Schindler *et al.* (1980) who argued LPS was highly mobile based on fluorescence recovery after photobleaching (FRAP) experiments measuring the mobility of exogenous, rhodamine-modified LPS molecules after their apparent integration into the OM of *S. typhimurium* G30A ^16^. This was disputed by Ghosh *et al*. (2005), who also used FRAP to monitor the average mobility of fluorescent concanavalin A labelled smooth LPS in the OM of *Escherichia coli* 2443 ^17^. The lack of consensus in these landmark studies likely arises from limitations associated with the LPS labelling methods, with physically large or multivalent exogenous fluorescent probes unsuitable for accurate *in vivo* characterisation as their binding to LPS potentially influences its mobility and organisation in the OM.

To overcome these challenges, we implement metabolic LPS engineering ^18,19^ to enable direct, *in situ* fluorescent labelling of the Lipid A-(Kdo)_2_ conserved within the LPS inner core (Figure 1D and Supplementary Figure 1). This specific and efficient bio-orthogonal labelling strategy allowed us to employ ensemble and single-molecule fluorescence microscopy to demonstrate that LPS is laterally confined. By characterising the molecular basis for this restriction, we provide new insight into the mechanisms that underpin the vital barrier and load bearing functions of the OM in Gram-negative bacteria ^9^.

## Results

### LPS is tightly confined in the OM irrespective of oligosaccharide domain size

In a two-step approach, bacteria were initially cultured with 3-deoxy-D-manno-oct-2-ulosonic acid (Kdo) monosaccharide modified with a bio-orthogonal azide group (Kdo-N_3_). Kdo-N_3_ was taken up via a salvage pathway, biosynthetically incorporated into the inner core oligosaccharide domain of LPS (Figure 1D and Supplementary Figure 1) ^18,19^ and then ‘click’-labelled at the cell surface via Cu(I)-catalysed (CuAAC) or Cu(I)-free strain promoted azide-alkyne cycloaddition (SPAAC) ^20,21^ with an exogenously delivered alkyne functionalised, small (<1000 Da) photostable fluorophore (*i.e*. AF488, AF568 and AZ647)(Supplementary Figures 2, 3 and 4). Using this site-specific bio-orthogonal labelling approach and FRAP, we unequivocally show that LPS is laterally confined in the outer leaflet of the OM of the *E. coli* K12 cell envelope (Figure 1B,E). Next, we compared the mobile fraction distributions of fluorescently labelled rough LPS (AF488-LPS) in the OM of intact *E. coli* K12 MG1655 cells with those of fluorescently labelled polymeric peptidoglycan (PG) in the cell wall ^22,23^. Both species exhibited very restricted average lateral mobility (Figure 1A-1C), and despite the difference in their molecular weights (average M_w_,_PG_ ≈ 3.9 × 10^9^ Da vs. M_w,rough LPS_ ≈ 3000 Da) no statistically significant difference was observed when FRAP-derived mobile fraction distributions were compared (*p* = 0.63, Supplemenatry Table 1). The immobility of LPS in the OM and PG in the cell wall sharply contrasted with that of a recombinant N-terminal GFP-tagged IM protein (GFP-TolA), shown previously to undergo Brownian diffusion (Figure 1A-1C) ^24^, with the lateral mobility of AF488-LPS significantly restricted compared to that of GFP-TolA (*p* = 4.4 × 10^−7^, Supplementary Table 1).

Having demonstrated LPS confinement in the context of the Gram-negative OM, we sought to identify and characterise the underlying intermolecular interactions. We began by investigating the influence of LPS oligosaccharide domain structure (Figure 1D) on lateral mobility, conducting FRAP experiments on labelled LPS in the OM of an *E. coli* K12 Δ*waaC* mutant strain capable of only producing deep rough LPS, *E. coli* MG1655 where rough LPS predominates, and an O-antigen restored (OAR) *E. coli* strain producing smooth LPS (Figure 1E) ^25,26^. Despite pronounced differences in the predominating LPS oligosaccharide domain structures (Figure 1D), inter-strain variations in LPS mobile fraction distributions were statistically insignificant (Supplementary Table 1), and LPS lateral mobility in all strains was tightly confined (Figure 1F).

### LPS assembles into supramolecular structures in the OM

We then employed two-colour direct stochastic optical reconstruction microscopy (dSTORM) to map the distributions of the fluorescently labelled LPS relative to an abundant OMP (OmpA, 10^5^ copies in the OM ^4^) found in OMP islands across *E. coli* cell surfaces ^27^ (Figure 2A and Supplementary Figure 5). Simultaneous OMP fluorescent-labelling was achieved using amber codon suppression ^28^, incorporating an alkyne functional group within an extracellular loop of OmpA (OmpA*) enabling azide-functionalised dye conjugation (Supplementary Figure 5). Interrogation of dSTORM data revealed distinct LPS-rich and OMP-rich regions within OMs that were independent of dye selection (Supplementary Figure 5), with median diameters of ∼0.26 µm and ∼0.18 µm, respectively (Figure 2B). Subsequent co-localisation analysis confirmed that LPS prefers self-assembly into supramolecular structures or islands (Figure 2C), as recently hypothesised ^27^. This discrete clustering of fluorescently labelled LPS allowed us to explore the factors influencing lateral mobility within these glycolipid-rich islands, thus facilitating dissection of dominant intermolecular LPS-LPS interactions over LPS-OMP interactions.

**Figure 2:**
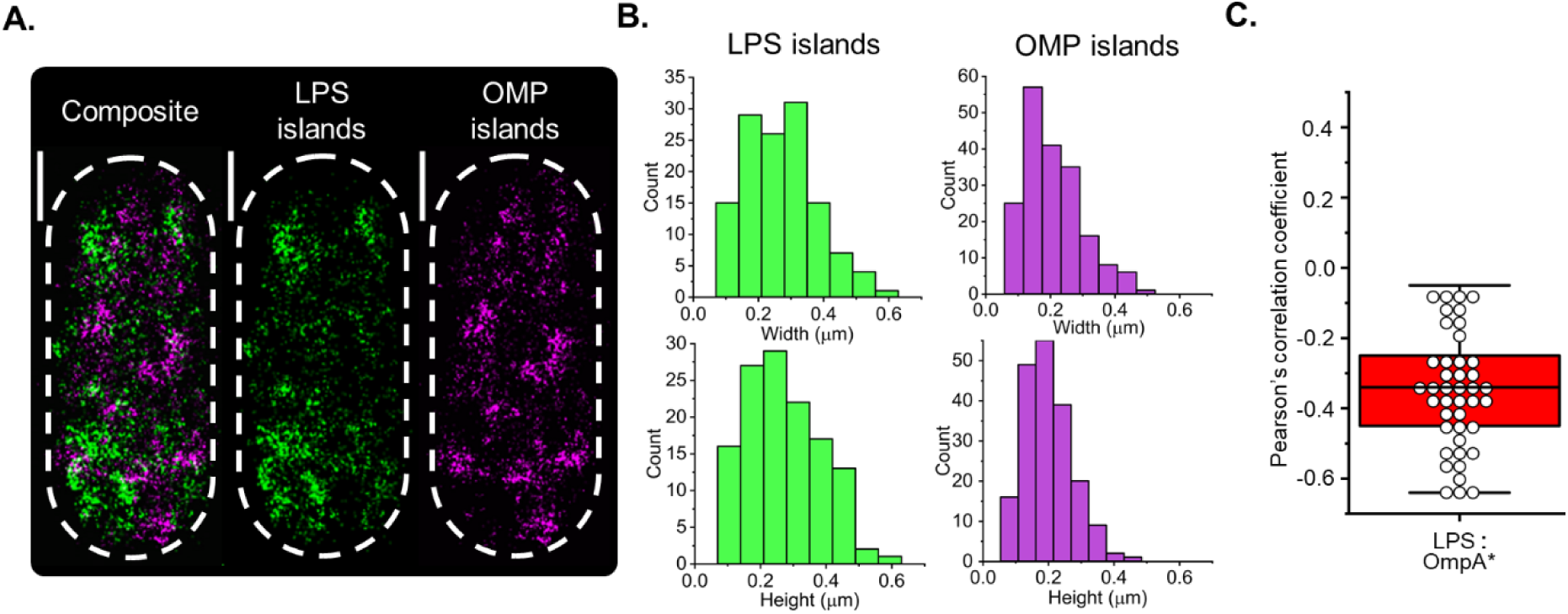
Two-colour direct stochastic optical reconstruction microscopy (dSTORM) reveals distinct LPS- and OMP-rich regions across the *E. coli* cell surface. **A.** Representative two-colour dSTORM images acquired using total internal reflection illumination showing the discrete clustering of AZ488-labelled LPS and AZ647-labelled recombinant non-canonical amino acid (ncAA) containing OmpA* in the OM of an *E. coli* Δ*ompA* cell. The discrete clustering of these abundant OM components, which is apparent in the composite image, indicates that LPS molecules and OMPs can assemble into their own spatially separated supramolecular islands in the OM. Scale bars: 0.5 µm. **B.** An automated thresholding algorithm was used to identify LPS- and OMP-rich regions or islands in the respective fluorescence channels, and the height (longest axis) and width (shortest axis perpendicular to long axis) were measured manually for all patches > 50 µm in size. Histograms of the width and height measurements for individual LPS islands (n = 127 islands from 31 cells) and OMP islands (n = 190 islands from 30 cells) indicated a broad range of island sizes existed in the OM. The median size of the LPS islands (median width: 0.260 ± 0.010 µm, median height: 0.267 ± 0.009 µm) was slightly greater than the OMP islands (median width: 0.189 ± 0.006 µm, median height: 0.180 ± 0.006 µm). All experiments were done in triplicate. **C.** Co-localisation analysis of two-colour dSTORM data confirmed the strong negative correlation (median Pearson correlation coefficient = - 0.35) between the surface location of most fluorescently labelled LPS and OmpA* in *E. coli* Δ*ompA* cells. All experiments were done in triplicate (n = 42).

### LPS lateral confinement is influenced by OM asymmetry and OM-PG covalent linkages

Accordingly, we investigated whether OM composition, asymmetry and covalent OM-PG associations influence average LPS lateral mobility. We compared the mobility of LPS in the MG1655 strain with that of LPS in the *imp4213* strain ^29^, in which OM asymmetry is compromised. In *imp4213* cells, an in-frame deletion in *lptD* disrupts LptD / LptE lipoprotein complex formation, reducing the rate of LPS insertion in the OM ^29^. The resulting decrease in LPS OM outer leaflet density prevents tight packing of adjacent LPS molecules, reducing the capacity for LPS-LPS interactions and creating spaces in the outer leaflet that are spontaneously filled with phospholipid (PL) from the inner leaflet. Our bio-orthogonal labelling approach enabled the low-level, site-specific fluorescent labelling of LPS required for single-molecule detection and localisation. Therefore, single-particle tracking (SPT) was used to directly probe whether sub-populations of LPS molecules with different lateral mobilities existed in the *imp4213* strain.

SPT of video data for AF488-LPS in the *imp4213* strain revealed two populations of rough LPS molecules with distinct diffusive characteristics (Figure 3A,C). One-third of the LPS molecules were observed to diffuse freely in the OM (D_2D_ = 0.0550 ± 0.00744 µm^2^/s), while two-thirds of the LPS molecules displayed restricted lateral diffusion (D_2D_ = 0.0347 ±0.00237 µm^2^/s) (Figure 3B and Supplementary Table 2). The lateral diffusion coefficients obtained for confined rough LPS (MG1655, 0.0182 ±0.000645 µm^2^/s) and confined deep rough LPS (Δ*waaC*, 0.0181 ±0.000727 µm^2^/s) were significantly slower in comparison and nearly identical to each other in magnitude (Supplementary Table 2). However, the confinement diameter estimated from the asymptotic mean-squared displacement (MSD) value was greater for the *imp4213* strain (0.723 ±0.0207 µm) relative to the MG1655 (0.543 ±0.00666 µm) and Δ*waaC* (0.543 ±0.00619 µm) strains (Supplementary Table 2). These observations suggest an overall reduction in LPS corralling within the supramolecular LPS islands (Figure 2), and a parallel increase in the lateral diffusion rate due to increased fluidity when more PL is present in the outer leaflet of the OM.

**Figure 3:**
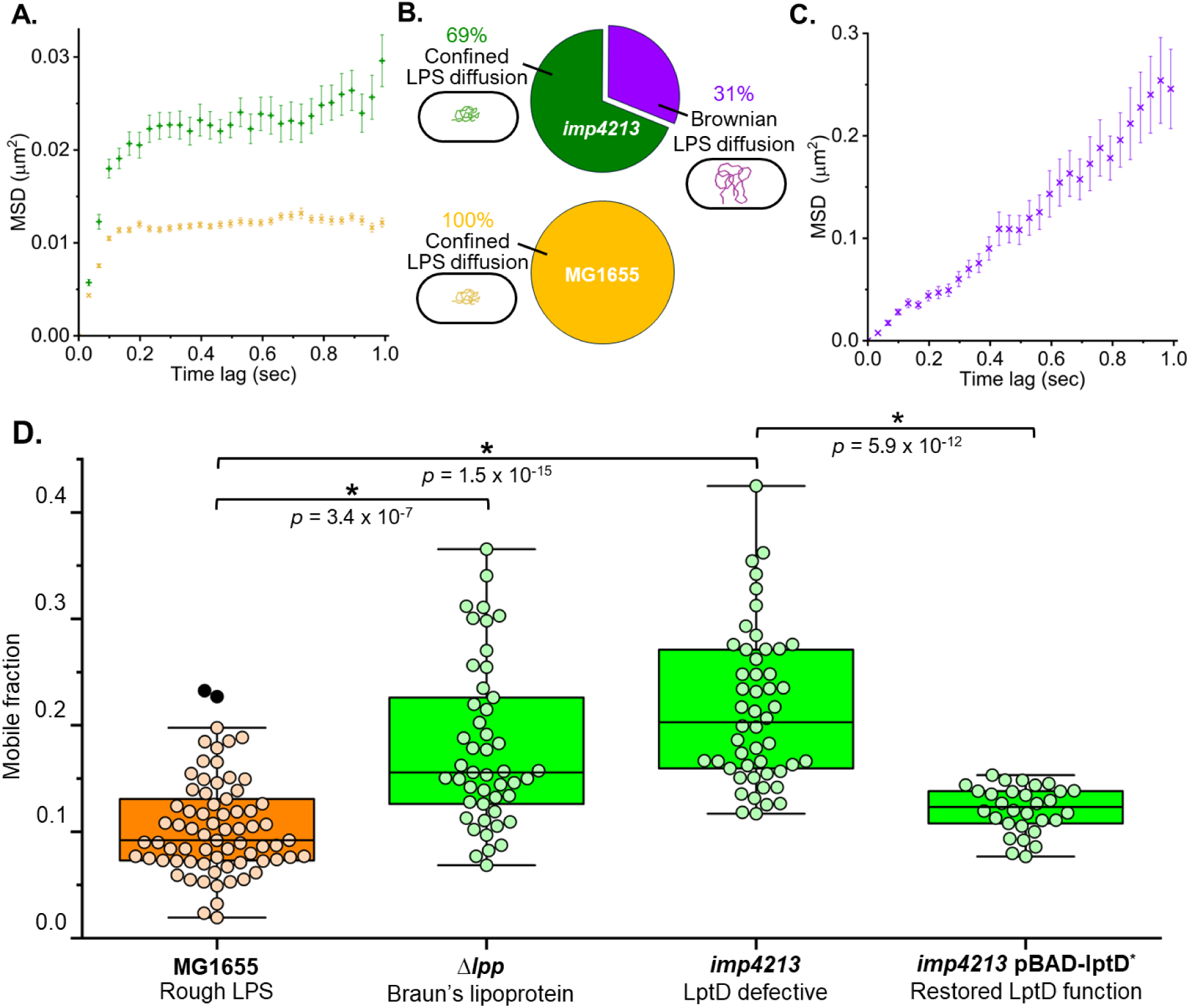
OM asymmetry, LPS-LPS intermolecular interactions, and OM-PG associations contribute to LPS confinement in the OM of *E. coli*. **A.** Mean-squared displacement (MSD) was calculated for single AF488-labelled LPS molecules that could be tracked in the OM for at least 1 s before photobleaching (error is reported as S.E.M.). TIRFM video data was collected at 30 Hz from a minimum of three experimental replicates. The MSD for all rough LPS in *E. coli* MG1655 cells (orange **×**, n = 156) and most rough LPS in *E. coli imp4213* cells (green **+**, n = 104) rapidly approached an asymptotic value which was consistent with confined lateral diffusion. Linear regression of the MSD for the first 4 time delays yielded lateral diffusion coefficients (D_2D_) of ∼0.0182 µm^2^/s and ∼0.0347 µm^2^/s, respectively, for rough LPS in these cells. **B.** The majority of rough LPS molecules (69%) tracked on *E. coli imp4213* cells displayed confined lateral diffusion. The remainder of the tracked LPS molecules (31%) displayed free Brownian lateral diffusion. All LPS molecules tracked on wild-type *E. coli* MG1655 cells displayed confined lateral diffusion. **C.** The MSD for freely diffusing rough LPS in *E. coli imp4213* cells (violet **×**, n = 49) increased linearly with time, and linear regression of the first 13 time delays yielded D_2D_ ≈ 0.0550 µm^2^/s. **D.** FRAP-derived mobile fractions for rough LPS in the OM of *E. coli* MG1655, Δ*lpp* and LptD defective *imp4213* cells were compared to *E. coli imp4213* pBAD-lptD* cells 2 h post-initiation of functional LptD* production. Statistically significant increases in LPS mobile fractions (by Mann-Whitney test) were observed in Δ*lpp* (median = 0.16, n = 47) and *imp4213* (median = 0.20, n = 50) cells relative to wild-type *E. coli* MG1655 cells (median = 0.09, n = 74). Restoration of OM asymmetry via production of functional recombinant LptD* in *imp4213* pBAD-lptD* cells (median = 0.12, n = 30) triggered a reduction in LPS lateral mobility to near wild-type levels.

Consistent with these single-molecule level observations, the LPS mobile fraction in the *imp4213* strain determined by FRAP was also significantly higher compared to the MG1655 strain (Figure 3D and Supplementary Table 1). These complementary data suggest that strict maintenance of OM asymmetry and tight, ordered packing of adjacent molecules to maximise the number and strength of LPS-LPS interactions is a pre-requisite for restricted lateral mobility of LPS in the OM. To reinforce this hypothesis, we demonstrated that restoration of OM asymmetry in the *imp4213* strain, through expression of a functional LptD protein capable of LptD / LptE complex formation (Supplementary Figure 6), resulted in a statistically significant decrease in the average LPS lateral mobility to near wild-type levels (Figure 3D and Supplementary Table 1). Rescued *imp4213* cells expressing functional LptD also showed reduced sensitivity to SDS, EDTA and bile salts (Supplementary Figure 6) demonstrating the links between OM asymmetry, LPS confinement and the barrier function of the OM.

The periplasmic surface of the OM is held near the bacterial cell wall by the very abundant Braun’s lipoprotein (Lpp) which is both covalently coupled to peptidoglycan and anchored in the membrane ^30^. We also measured the average lateral mobility of LPS in an *E. coli* Δ*lpp* strain, shown previously to have decreased OM stiffness and stability ^31,32^. A statistically significant increase in the lateral mobility of LPS was observed in the Δ*lpp* strain (*p* = 3.7 × 10^−7^, Supplementary Table 1) relative to the MG1655 strain (Figure 3D). Previously, we showed that OMP lateral mobility was not affected by the deletion of *lpp* ^33^. This implies that the separate LPS-rich and OMP-rich regions of the OM may respond differently to the decrease in OM stiffness and stability present in the Δ*lpp* strain. Comparable increases in the lateral mobility of LPS were observed in the Δ*lpp* and *imp4213* strains (Figure 3D), even though the origin of the OM perturbations is very different in these strains. Since the majority of LPS remained immobile in these mutants (Figure 3D), it is possible that the supramolecular assemblies in the OM incorporate redundant biophysical interactions that organise and stabilise them.

### Divalent cation-mediated interactions influence LPS mobility

LPS is anionic glycolipid with a formal charge density of up to 1.0 per hydrocarbon chain ^5^, which is greater than PL where formal negative charge density per hydrocarbon chain is 0.5 ^5^. Divalent cation (Ca^2+^ and Mg^2+^) coordination by negatively charged substituents (PO_4_^2−^ and CO_2_^−^) within the conserved Lipid A-(Kdo)_2_ core of LPS have been proposed to neutralise these potentially repulsive charges enabling tighter LPS packing in the OM ^13,34^ and are therefore considered critical for maintenance of OM asymmetry, stability, and integrity ^35^. However, our current understanding of the affinity of LPS molecules for specific divalent cations and the resulting influence on ionic bridge formation between adjacent anionic oligosaccharide head groups in the OM relies heavily on *in silico* modelling ^14,35–38^. For instance, a recent computational study concluded that Ca^2+^ (rather than Mg^2+^) mediates interactions between the PO_4_^2−^ groups of β-1,6-glucosamine disaccharides in Lipid A and exerts the greatest stabilising influence on LPS in the OM ^35^. But current models are limited by the complexities of LPS, particularly the chemical heterogeneity of smooth phenotypes consisting of O-antigen-containing LPS of different lengths. This is significant because most of the Gram-negative bacteria that exist outside the laboratory — such as those that occupy pathological niches — produce O-antigen-containing smooth LPS as a key virulence factor ^39^.

Since FRAP results (Figure 1E,F) demonstrated deep rough, rough and smooth LPS were all tightly confined in the OM, we reasoned that mobility defining interactions must involve groups common to all these LPS glycoforms. This potentially includes divalent cation-mediated interactions within the conserved, negatively charged Lipid A-(Kdo)_2_ motif at the base of LPS. To explore this hypothesis, we incubated cells presenting deep rough, rough and smooth fluorescent LPS with divalent cation chelating agents EDTA and EGTA prior to FRAP analysis. Although both chelating agents co-ordinate Mg^2+^ and Ca^2+^, EGTA has a five-fold higher affinity for Ca^2+^ compared to Mg^2+^ while EDTA has approximately equivalent affinities for both Mg^2+^ and Ca^2+ 40,41^. Comparison of the impacts of EDTA versus EGTA on average LPS lateral mobility therefore provided insights into the relative influence of Mg^2+^ (EDTA treatment) and Ca^2+^ (EGTA treatment) mediated interactions on LPS confinement. Repeating experiments with strains producing deep rough (Δ*waaC*), rough (MG1655) or smooth LPS (OAR) provided insight into the relative importance of divalent cation mediated interactions compared to other potential forms of intermolecular interactions (such as hydrogen bonds) likely to exist between the oligosaccharide domains of adjacent LPS molecules. These FRAP experiments confirmed that divalent cation mediated interactions between adjacent LPS molecules do contribute toward LPS lateral restriction in *E. coli*, with statistically significant positive shifts in LPS mobile distributions observed for all *E. coli* strains after EDTA or EGTA treatments (Figure 4A,B and Supplementary Table 1). However, smooth LPS lateral mobilities were less susceptible to chelator treatment which suggests the extended hydrogen bond networks in the LPS O-antigen repeats may offset the destabilising impact of divalent cation removal, and / or increase steric hindrance limiting the efficacy of chelator treatment. Interestingly the increase in LPS mobility upon chelator treatment was not replicated with *imp4213* cells (Supplementary Figure 7 and Supplementary Table 1). The apparent insensitivity of *imp4213* cells to chelator treatments suggests that loss of OM asymmetry and tight packing of adjacent LPS molecules also contributes to disruption of divalent cation-mediated LPS-LPS interactions.

**Figure 4:**
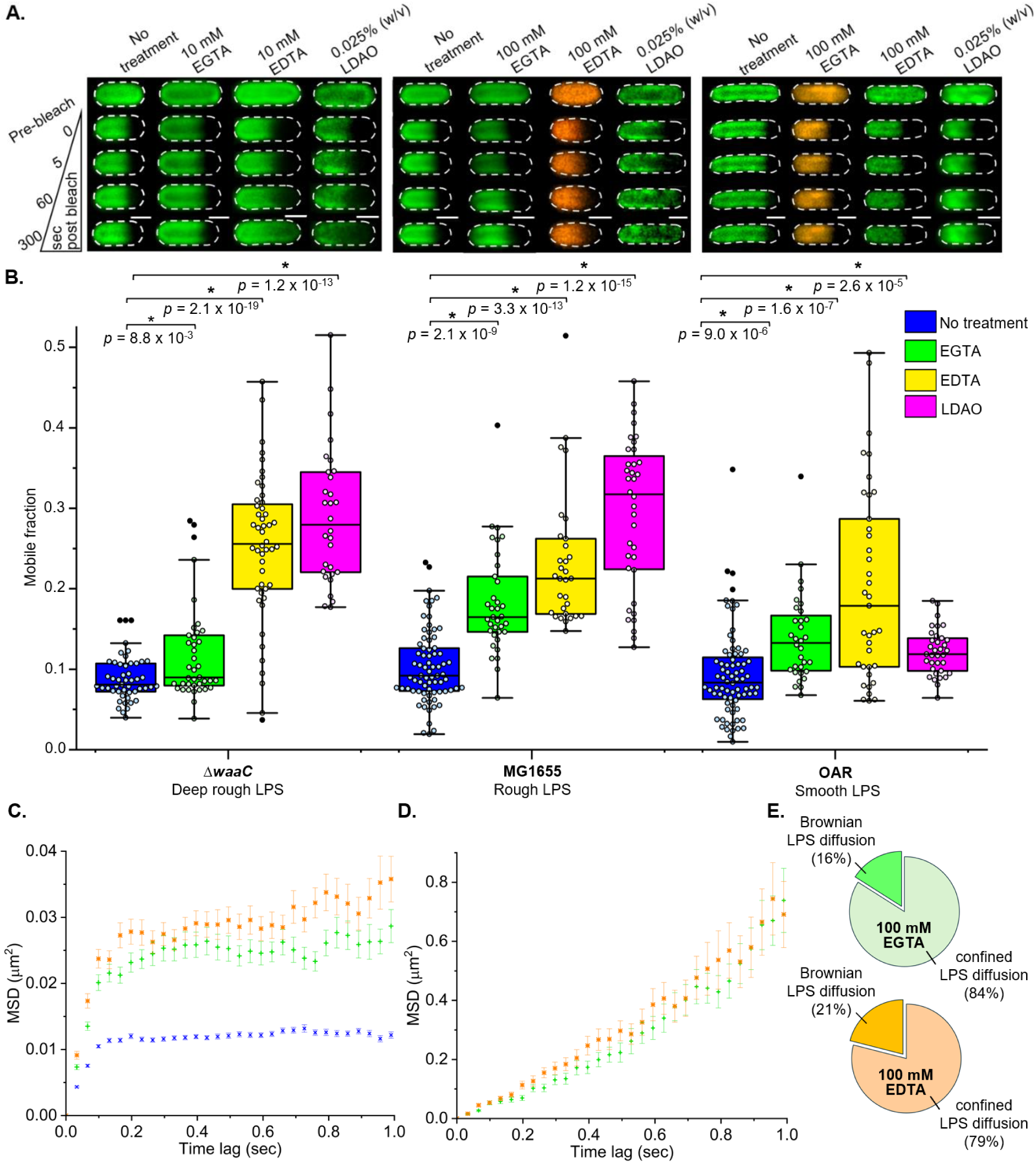
Confined lateral diffusion of LPS in the OM of *E. coli* is influenced by divalent cation-mediated interactions and hydrophobic interactions within the membrane. **A.** Representative FRAP sequences for AF488-LPS (green) or AF594-LPS (orange) in the OM of deep rough LPS-producing *ΔwaaC* (left), rough LPS-producing MG1655 (middle), and smooth LPS-producing OAR (right) *E. coli* cells without treatment, and after treatment with chelator (EGTA or EDTA) or detergent (0.025% (w/v) LDAO). Scale bars: 1.0 µm. **B.** Comparison of FRAP-derived LPS mobile fractions in *E. coli* cells producing different LPS glycoforms: *ΔwaaC* (untreated, n = 49; 10 mM EGTA, n = 39; 10 mM EDTA, n = 48; LDAO, n = 30), MG1655 (untreated, n = 74; 100 mM EGTA, n = 35; 100 mM EDTA, n = 31; LDAO, n = 37) and OAR (untreated, n = 75; 100 mM EGTA, n = 32; 100 mM EDTA, n = 37; LDAO, n = 35). **C.** Collection of TIRFM video data, and the calculation and analysis of the MSD (± S.E.M.) for single AF488-labelled LPS molecules in the OM of *E. coli* MG1655 cells was done as described in Figure 3. The MSD for all rough LPS in untreated cells (blue **×**, n = 156) rapidly approached an asymptotic value which was consistent with confined lateral diffusion. The majority of rough LPS molecules tracked on 100 mM EDTA (orange **®**, n = 118) and 100 mM EGTA (green **+**, n = 108) treated cells also had an MSD (≈ 0.03 µm^2^) which was consistent with confined lateral diffusion, but the degree of confinement was reduced relative to untreated cells (≈ 0.01 µm^2^). Linear regression of the MSD yielded D_2D_ ≈ 0.0377 µm^2^/s and D_2D_ ≈ 0.0373 µm^2^/s for confined rough LPS in EDTA- and EGTA-treated cells, respectively. These values were increased relative to untreated cells (D_2D_ ≈ 0.0182 µm^2^/s). **D.** The MSD for some rough LPS molecules tracked on 100 mM EDTA-treated (orange **®**, n = 31) and 100 mM EGTA-treated (green **+**, n = 21) cells increased linearly with time which indicated free Brownian diffusion. Linear regression yielded D_2D_ ≈ 0.153 µm^2^/s and D_2D_ ≈ 0.113 µm^2^/s, respectively, for these cells. **E.** Percentage of rough LPS molecules observed to undergo confined and free Brownian lateral diffusion in the OM of 100 mM EDTA-treated and 100 mM EGTA-treated cells.

SPT of video data for fluorescently-labelled LPS in the MG1655 strain after treatment with EDTA or EGTA revealed two populations of LPS molecules with distinct diffusive characteristics (Figure 4C,E). The majority of the LPS species remained confined when treated with chelator (79% for EDTA, 84% for EGTA); however, a new population of freely diffusing LPS molecules was also observed (21% for EDTA, 16% for EGTA) (Figure 4E). The lateral diffusion coefficient of the confined LPS species were similar for both chelators (EDTA: D_2D_ = 0.0377 µm^2^/s; EGTA: D_2D_ = 0.0373 µm^2^/s) and slightly increased relative to untreated cells (Figure 4D and Supplementary Table 2). The confinement diameter for LPS also increased in the MG1655 strain upon chelator treatment (EDTA: 0.815 ±0.0238 µm; EGTA: 0.748 ±0.0200 µm) yielding values equivalent to what was observed for confined diffusion in the *imp4213* strain (Supplementary Table 2). In our previous study, a fluorescent OMP-labelling strategy was used to follow the lateral diffusion of single CirA receptors (a monomeric OMP) in the bacterial OM by SPT ^33^, and it was used here to probe how chelator treatment affected lateral diffusion in OMP-rich islands. In striking contrast to LPS, all of the CirA receptor displayed confined diffusion in the presence of these chelators (Supplementary Figure 8) with the lateral diffusion coefficient (EDTA: D_2D_ = 0.0454 ±0.00340 µm^2^/s; EGTA: D_2D_ = 0.0402 ±0.00471 µm^2^/s) and confinement diameter (EDTA: 0.846 ±0.0229 µm; EGTA: 0.740 ±0.0238 µm) increased marginally relative to untreated cells (D_2D_ = 0.0178 ±0.000623 µm^2^/s and 0.540 ±0.00665 µm, respectively)(Supplementary Table 2). These SPT results are again consistent with divalent cation depletion in the OM disrupting LPS-LPS interactions specifically while LPS-OMP interactions remain mostly intact.

### Mg^2+^ rather than Ca^2+^ ions primarily influence LPS lateral mobility

Notably, EDTA treatment triggered larger shifts in LPS mobile fractions compared to EGTA treatment (Figure 4 and Supplementary Table 1) implying that Mg^2+^ mediated interactions exert a greater restrictive influence on LPS diffusion compared to Ca^2+^. This contrasts with the conclusions of computational studies that concluded Ca^2+^-mediated interactions are solely responsible for LPS OM confinement ^14,42^. Alternatively, it may be that EDTA is more effective at stripping OM-coordinated divalent cations compared to EGTA. Categorical determination of the relative influence of Mg^2+^ versus Ca^2+^-mediated OM interactions on LPS average mobility from these FRAP experiments alone is difficult because both EGTA and EDTA will chelate Ca^2+^ and Mg^2+^. Analyses are further complicated by the composition of the standard chemically defined medium (CDM) used in this work which contains 2.0 mM Mg^2+^ and 0.1 mM Ca^2+^. Therefore, we conducted a series of experiments growing cells in CDM with altered Mg^2+^ and Ca^2+^ ion ratios after verification that these changes did not affect cell morphologies or growth rates. We observed that LPS mobile fraction distributions in the OM of cells cultured in low Mg^2+^ (0.1 mM), low Ca^2+^ (0.1 mM) CDM were significantly elevated compared with those measured in cells cultured in standard, high Mg^2+^, low Ca^2+^ CDM (Figure 5, Supplementary Tables 1 and 3). Restoration to original levels of LPS restriction in standard CDM (with high Mg^2+^ and low Ca^2+^) were not observed when the Ca^2+^ concentration was increased to 2.0 mM. These results support our initial hypothesis that Mg^2+^ mediated interactions exert a greater restrictive influence on LPS lateral diffusion compared to Ca^2+^ interactions, and reinforce the conclusions drawn from chelator treatment experiments (Figure 4).

**Figure 5:**
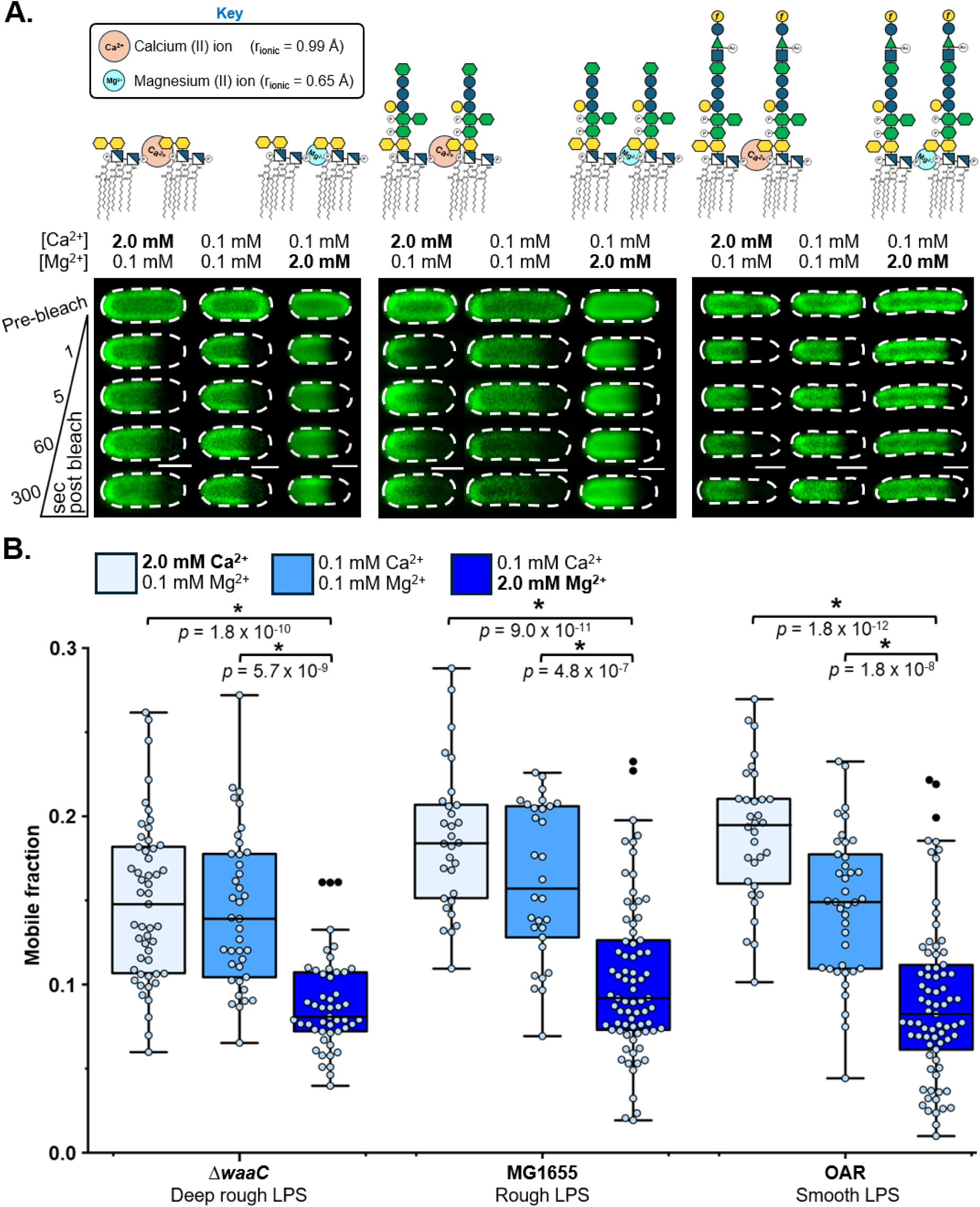
A reduction in the Mg^2+^ ion concentration triggers an increase in LPS lateral mobility in the OM of *E. coli* cells. **A.** Schematics of deep rough, rough and smooth LPS glycoforms present in Δ*waaC*, MG1655 and O-antigen restored (OAR) *E. coli* strains, respectively, showing how calcium (II) and magnesium (II) ions can bridge adjacent LPS molecules by binding to phosphate groups within their conserved Lipid A cores. The smaller physical size (ionic radius = 0.65 Å) and higher charge density of the Mg^2+^ ion will permit tighter packing and stronger interactions between adjacent LPS molecules in the OM compared to the physically larger Ca^2+^ ion (ionic radius = 0.99 Å) with a lower charge density. Representative FRAP sequences for AF488-LPS in the OM of deep rough LPS-producing *ΔwaaC* (left), rough LPS-producing MG1655 (middle), and smooth LPS-producing OAR (right) *E. coli* cells after culturing in chemically defined medium (CDM) with varied Mg^2+^ and Ca^2+^ ion concentrations at constant total divalent cation concentration. Scale bars: 1.0 µm**. B.** Comparison of FRAP-derived LPS mobile fractions in *E. coli* cells producing different LPS glycoforms after culturing in CDM with varied Mg^2+^ and Ca^2+^ ion concentrations. Bacterial cells cultured in high Ca^2+^ (2.0 mM), low Mg^2+^ (0.1 mM) CDM or low Ca^2+^ (0.1 mM), low Mg^2+^ (0.1 mM) CDM displayed a statistically significant increase in the median LPS mobile fraction (by Mann-Whitney test) compared to cells cultured in low Ca^2+^, high Mg^2+^ CDM, irrespective of the LPS glycoform produced by the *E. coli* strain. This demonstrates that Mg^2+^ mediated LPS-LPS interactions exert a greater restrictive influence on LPS lateral mobility in the OM compared to Ca^2+^ mediated interactions.

### Hydrophobic forces within the OM contribute to restricted LPS lateral mobility

Even with chelator treatment or reduced Mg^2+^ concentration the majority of LPS molecules in the OM outer leaflet remain immobile irrespective of LPS oligosaccharide domain size (Figures 4 and 5). Under standard conditions most species of Gram-negative bacteria produce hexa-acylated Lipid A in LPS (Supplementary Figure 1), with all six acyl chains being saturated, facilitating tight packing of adjacent LPS molecules whilst providing a large potential area for hydrophobic interactions (*e.g*. van der Waals contacts). We therefore theorised these interactions might also contribute to our observed high levels of LPS restriction. To investigate we subjected cells to detergent treatment at a sub-lytic concentration (Figure 4A,B). *N,N*-Dimethyldodecylamine N-oxide (LDAO) is a zwitterionic, OM-targeting detergent used for selective isolation of OMPs with the ability to trigger changes in LPS monolayer structure ^43,44^. We observed that pre-FRAP treatment with 0.025% (w/v) LDAO triggered positive shifts in the LPS mobile fraction distributions of all three *E. coli* strains (Figure 4A,B) confirming that interactions within the hydrophobic environment of the OM do contribute to LPS restriction. As observed in FRAP experiments after chelator treatment, the size of positive mobility shifts was influenced by LPS oligosaccharide domain structure, with larger increases observed in cells with deep rough and / or rough LPS, highlighting the steric protective effect the extended O-antigen affords to the conserved Lipid A-(Kdo)_2_ motif of LPS. Restrictive O-antigen-mediated intermolecular hydrogen bonds in the OM, unlikely to be influenced by detergent treatment, may also offset the destabilising impact of LDAO molecules that do penetrate through the O-antigen. We tested whether treatment with a non-lethal concentration of chaotropic agent (300 mM urea) disrupted the LPS oligosaccharide domain structure and increased lateral mobility, but only a minor effect on mobility was observed (Supplementary Figure 9). Our experimentally derived relationship between LPS oligosaccharide domain size and OM detergent susceptibility reinforces previous studies that have emphasised the importance of LPS O-antigens in conveying enhanced resistance to bile salts in the context of GI tract infections ^45^.

The increase in lateral mobility of LPS in the deep rough (Δ*waaC*) strain induced by 0.025% (w/v) LDAO treatment is consistent with the detergent molecule disrupting the optimal hydrophobic packing between LPS molecules. As LDAO is used to solubilise OMPs ^43,44,46^, the increases in LPS mobility may also result from the preferential disruption of LPS-OMP interactions rather than LPS-LPS interactions. We therefore utilised an endotoxin-free *E. coli* strain (ClearColi™) ^47^ which incorporated tetra-acylated Lipid IV_A_ (Supplementary Figure 10A), rather than hexa-acylated Lipid IV_A_, to dissect whether the detergent was indeed disrupting hydrophobic LPS-LPS interactions. Once metabolically labelled with Kdo-azide, the tetra-acylated Lipid IV_A_ (termed LPS_4-acyl_) has an oligosaccharide headgroup (two Kdo sugars) equivalent to the native deep rough LPS (Supplementary Figure 10A). However, we hypothesised that the LPS_4-acyl_ would prevent optimal acyl chain packing and increase lateral mobility. We observed exactly this result for the LPS_4-acyl_ strain in the standard CDM by FRAP (Supplementary Figure 10B,C). The observed LPS_4-acyl_ lateral mobility in the ClearColi™ strain was like that observed in the deep rough (Δ*waaC*) strain after 0.025% (w/v) LDAO treatment (Figure 4A,B). These results are consistent with an increase in the lateral mobility of LPS due to sub-optimal acyl chain packing. To ascertain whether the OMP-rich regions of the OM were also perturbed in the ClearColi™ strain, we used fluorescent labelling to track the lateral diffusion of the CirA receptor with SPT in the ClearColi™ strain, before and after metabolic labelling with Kdo sugar (Supplementary Figure 10D). The observed lateral diffusion coefficients (D_2D_ = 0.0181 ±0.000820 µm^2^/s and D_2D_ = 0.0229 ±0.00206 µm^2^/s) and confinement diameters (0.546 ±0.00760 µm and 0.600 ±0.0124 µm) of this OMP in the ClearColi™ strain before and after metabolic labelling with Kdo sugar were nearly indistinguishable from that observed for the MG1655 strain (Supplementary Figure 10D and Supplementary Table 2). This emphasises the importance of Lipid A acyl chain number in determining the fluidity of the LPS-rich OM regions, while intimating that the packing of LPS molecules within OMP-rich regions may be different and highly influenced by LPS-OMP interactions.

### LPS confinement is conserved in pathogenic Gram-negative bacteria

Although the general structure of Lipid A and inner core oligosaccharides is highly conserved across Gram-negative bacteria, significant interspecies differences in outer core and O-antigen repeat structure exist ^48–50^. Having demonstrated that, under chemical challenge, differences in *E. coli* K12 LPS oligosaccharide domain structure influence average LPS mobility, we elected to carry out selected FRAP experiments on other Gram-negative bacteria. We characterised LPS mobility in *Salmonella enterica* serovar Typhimurium LT2, *Pseudomonas aeruginosa* PAO1 and *E. coli* UTI89 strains to ascertain the extent to which observations for *E. coli* K12 could be extrapolated across important pathogens within the Gram-negative clade. *P. aeruginosa* PAO1 and *E. coli* UTI89 exhibit a capsular serotype, a proven virulence factor. To enable fluorophore bioconjugation to Kdo-N_3_ in the LPS of these strains, an adapted labelling protocol was developed. This involved capsular polysaccharide stripping using EDTA followed by the restoration of original OM-coordinated divalent cation concentrations via ‘bathing’ of ‘stripped’ cells in freshly supplemented CDM, prior to fluorescent labelling. This method enabled comparison of average LPS mobilities and the impact of chelator treatments in bacteria with and without capsular serotypes.

The level of restricted lateral mobility for LPS was roughly equivalent in all strains, irrespective of whether cells had capsule-forming or non-capsule-forming serotypes (Figure 6, Supplementary Tables 1 and 4). These experiments reinforce the hypothesis that LPS confinement is a universal trait conserved across Gram-negative bacterial species with conventional OM physiology, including those that occupy pathological niches. Similarly, the restrictive influence of divalent cation mediated interactions in the OM on lateral LPS mobility is not a trait exclusive to *E. coli* K12 laboratory strains, as pre-FRAP treatment with 50 mM EDTA and 50 mM EGTA triggered a statistically significant increase in LPS mobility in all bacteria (Figure 6 and Supplementary Table 1). This further supports our assertion that the molecular basis of LPS confinement resides in the structure of Lipid A and the inner core oligosaccharide domain, and explains why these structural motifs are highly conserved across the Gram-negative bacterial clade ^39^.

**Figure 6:**
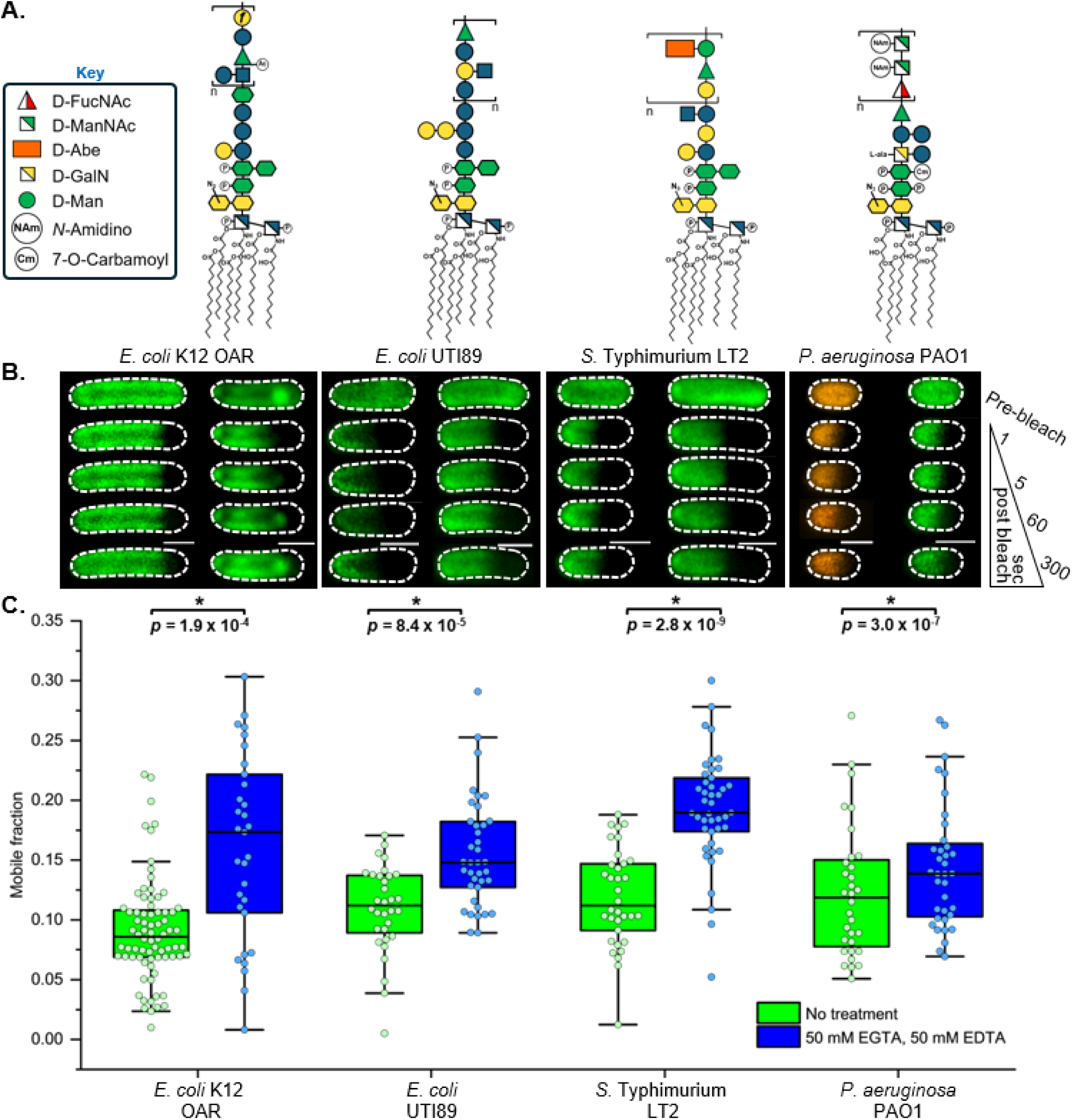
Divalent cation-dependent lateral confinement of LPS is observed in pathogenic Gram-negative bacteria, irrespective of O-antigen and capsule-forming serotypes. **A.** Schematics of smooth LPS glycoforms present in *E. coli* K12 O-antigen restored (OAR), *E. coli* UTI89, *S.* Typhimurium LT2 and *P. aeruginosa* PAO1 strains showing the composition of the oligosaccharide domain displayed at the cell surface. The *E. coli* UTI89 and *P. aeruginosa* PAO1 strains produce a capsular polysaccharide which was removed prior to fluorescent-labelling of the LPS. **B.** Representative FRAP sequences for AF488-LPS (green) or AF594-LPS (orange) in the OM of smooth LPS-producing strains before (left-hand image sequence) and after (right-hand image sequence) combined 50 mM EDTA and 50 mM EGTA treatment. The combined chelator treatment was found to be as effective as separate treatments with 100 mM EDTA and 100 mM EGTA at removing coordinated Mg^2+^ and Ca^2+^ ions from the OM. Scale bars: 1.0 µm. **C.** Comparison of FRAP-derived LPS mobile fractions for fluorescently labelled-LPS in the OM of *E. coli* K12 OAR (n_pre-treatment_ = 75, n_post-treatment_ = 31), *E. coli* UTI89 (n_pre-treatment_ = 32, n_post-treatment_ = 38), *S.* Typhimurium LT2 (n_pre-treatment_ = 34, n_post-treatment_ = 43) and *P. aeruginosa* PAO1 (n_pre-treatment_ = 30, n_post-treatment_ = 30) cells pre- and post-treatment with the combined chelators. LPS lateral mobility is tightly confined in the *E. coli* UTI89, *S.* Typhimurium LT2 and *P. aeruginosa* PAO1 pathogenic strains. The combined chelator treatment triggered a statistically significant increase in the lateral mobility of the fluorescently labelled LPS (by Mann-Whitney test) in the OM of each O-antigen producing strain, irrespective of the oligosaccharide domain composition and the degree of acylation in the Lipid A core.

## Discussion

Our characterisation of LPS lateral mobility using *in vivo* ensemble-averaged (FRAP) and single-molecule (SPT) fluorescence microscopy demonstrates unequivocally that this essential glycolipid is tightly restricted in native Gram-negative bacterial OMs. Remarkably, the observed confinement is not dependent on oligosaccharide headgroup size or structure (Figure 2), and this includes LPS from the K and O serotype groups (Figure 6). Rather, we identify bridging Mg^2+^ ions and saturated acyl chains at the base of LPS as principally responsible for LPS lateral confinement in the OM. The confinement diameters estimated from SPT data for deep rough and rough LPS, and for the CirA receptor with these glycoforms, were very similar (∼0.540 µm) (Supplementary Table 2) and are consistent with the formation of LPS- and OMP-rich regions as observed in our dSTORM data (Figure 2B). Our FRAP, SPT and super-resolution imaging data support the view that the OM is a mosaic of OMP and LPS-rich regions ^27^. OM confinement generates spatial heterogeneity at the bacterial cell surface, which governs how different OM components (and virulence factors) are inherited by daughter cells during binary fission. This creates cell-to-cell variation in a bacterial population which is known to influence the host immune response ^51^ and the success of antibiotic treatment ^52^.

OMP-rich regions in the OM are located in supramolecular islands ^33^, where each OMP is surrounded by a ring-like shell of asymmetric lipid which mediates interactions between neighbouring OMPs ^53^, thus stabilising the supramolecular structure. Our observations indicate that LPS molecules likely exist in two very different environments within the OM, each with potentially different lateral mobilities. In agreement with previous computational studies, we showed divalent cation mediated interactions involving charged groups at the base of the LPS inner core oligosaccharide domain exerted a significant influence on LPS lateral diffusion ^14,42^. However, in contrast to these studies, we show that Mg^2+^ mediated interactions exert a greater restrictive influence on LPS diffusion compared to Ca^2+^. Divalent cation depletion in the OM generates a sub-population of LPS molecules undergoing Brownian free diffusion, which is not normally present, while OMPs only experience a moderate increase in the lateral diffusion coefficient and confinement diameter determined using SPT. This suggests that the LPS-OMP interactions that exist in OMP islands are more resilient to chemical challenges compared to the LPS-LPS interactions that dominate in the LPS-rich regions, and that a global increase in LPS lateral mobility may signal a reduction in mechanical integrity of the OM.

The chemical structure and conformation of LPS facilitate the formation of multiple types of intermolecular LPS-LPS interactions, contributing to the tight packing of LPS molecules in the outer leaflet of the OM required for its aforementioned barrier and load bearing functions ^1,2,32^. It appears based on this experimental work that the optimal packing of acyl chains and the binding of Mg^2+^ are critical to maintaining native van der Waals contacts and electrostatic interactions between the Lipid A moieties of neighbouring LPS molecules. These observations also align with our understanding of how and why bacteria make chemical modifications to the Lipid A anchor of LPS that is already at the cell surface in order to alter the properties of the OM ^54^. The extent to which lateral diffusion in the OM is altered in response to external challenges or changes in membrane composition appears dictated by the different biophysical properties of LPS- and OMP-rich regions ^9^, with increases in LPS mobility an apparent indicator of decreases in OM stiffness and/or stability. The new insight gained by revealing the molecular basis of LPS confinement in the OM can assist in the design and development of novel OM-disrupting antimicrobial therapies and will improve our ability to build more accurate *in silico* models of the Gram-negative cell envelope.

## Methods

### Bacterial strains and plasmids

Details of strains used in this study are provided in Supplementary Table 5. Keio collection mutants of *Escherichia coli* K12 substr. BW25113 used in this work (Δ*waaC*, Δ*lpp,* Δ*ompA*) were validated previously via PCR, confirming the presence of kanamycin resistance cassette within the open-reading frame of interest ^25,33^. Plasmid pNP4 encoding GFP-TolA was kindly provided by Piet de Boer ^55^ and transformed into the *E. coli* K12 BW25113 strain ^56^. The in-frame deletion in the *lptD* gene was introduced into the *E. coli* K12 BW25113 strain using P1 transduction by Dr. Emmanuele Severi (University of York) using a previously published donor MC4100 strain with the *imp4213* allele ^29^. Introduction of this allele was verified by Sanger sequencing (Eurofins Genomics LLC) after its amplification by PCR (using wild-type *lptD* amplification primers, Supplementary Table 6). *Pseudomonas aeruginosa* PAO1 is a model pathogenic strain ^57^ and was obtained from Prof. Gavin Thomas (University of York). *Salmonella enterica enterica* serovar Typhimurium strain LT2 ^58^ (abbreviated to *S.* Typhimurium LT2 in this work) and *E. coli* UTI89 strains ^59^ were provided by Prof. Marjan van der Woude (University of York). The endotoxin-free ClearColi™ K12 strain was commercially available ^47^. The O-antigen restored (OAR) *E. coli* DFB1655 strain was obtained from Prof. Ian Henderson and Dr. Doug Browning (University of Birmingham) ^26^.

The pBAD-lptD* plasmid was produced via Gibson isothermal assembly ^60^. PCR was used to amplify the *lptD* gene insert (see Supplementary Table 6 for primers) from purified *E. coli* K12 subs. MG1655 genomic DNA. The amplified gene was cloned into a pBADcLIC vector using a commercial Gibson isothermal assembly kit (NEBuilder® HiFi DNA Assembly). PCR site-directed mutagenesis (see Supplementary Table 6 for primers) was then used to incorporate the amber stop codon TAG in place of the codon for the D600 residue (Supplementary Figure 6A). Successful cloning and mutagenesis were verified via DNA sequencing (Eurofins Genomics LLC). The pEVOL-pylRS plasmid, encoding the Pyrrolysyl-tRNA synthetase/pyrrolysyl-tRNA pair from *Methanosarcina mazei* was obtained from Dr. Edward Lemke (EMBL Heidelberg) under MTA ^28,61^. pBAD-lptD* and pEVOL-pylRS were co-transformed into *imp4213* via electroporation (MicroPulser, BioRad Laboritories) (1.8 kV, 25 µF, time constant: 3.5 ms). *Imp4213* pBAD-lptD^D600^ pEVOL-pylRS were selected by incubating co-transformed cells overnight at 37 °C on an Lysogeny Broth (Miller formulation, LB-Miller) agar plate containing 50 µg mL^−1^ ampicillin and 17.5 µg mL^−1^ chloramphenicol. Unless otherwise specified, the incubation of bacterial cultures was done at 37 °C with 220 rpm agitation in a 50 mL screw-cap polypropylene tube. Liquid pre-cultures were prepared by inoculating 5 mL supplemented M9 chemically defined medium (M9 CDM, Supplementary Table 7) with appropriate antibiotics (Supplementary Table 5) using a well-isolated single colony of the desired strain picked from freshly streaked LB-Miller agar plate. Post-inoculation cultures were incubated for 6 – 8 h or until cell densities reached a minimum optical density at 600 nm (OD_600_) of 1.5.

The pBAD-ompA* plasmid was produced using an analogous method to that employed in the production of pBADcLIC-lptD amber stop codon containing plasmids. Briefly, the *ompA* gene from *E. coli* K12 BW25113 was amplified via PCR from purified bacterial chromosomal DNA and integrated in the pBADcLIC vector via NEBuilder® HiFi DNA Assembly (see Supplementary Table 6 for primers). PCR-based site-directed mutagenesis was then used to mutate the codon for E89 to an amber stop codon (TAG) in the pBAD-ompA plasmid. Following plasmid purification (Qiagen© Midi plasmid purification kit) and DNA sequencing (Eurofins Genomics LLC), the TAG-containing pBAD-ompA vector (pBAD-ompA*) and pEVOL-pyIRS were co-transformed into *E. coli* Δ*ompA* cells via electroporation as above. Co-transformed cells were selected using LB-Miller agar plates with 30 µg mL^−1^ kanamycin, 100 µg mL^−1^ ampicillin and 35 µg mL^−1^ chloramphenicol. A single colony from this plate was used to inoculate a volume of supplemented M9 CDM with kanamycin, ampicillin and chloramphenicol antibiotics. Following prolonged pre-culture incubation, the appropriate volume of pre-culture was used to inoculate a fresh batch of supplemented M9 CDM with ampicillin and chloramphenicol antibiotics to an OD_600_ of 0.05. The cultures were then incubated for 2 – 3 hours.

### Colicin Ia purification and fluorophore-labelling

The colicin Ia probe was engineered with an inactivating disulfide (*top-lock*) in the coiled-coil receptor-binding domain (L257C, A411C) allowing high-affinity binding to the CirA receptor at the surface of *E. coli* cells but preventing translocation through the cell envelope and bacterial intoxication. The colicin Ia probe also had a solvent exposed cysteine in the cytotoxic domain (Cys544) for maleimide-directed fluorescent-labelling. The colicin Ia based probe used in this work was purified and fluorescently-labelled with the Alexa Fluor 488-maleimide (Invitrogen) as described previously ^33^. To increase the labelling efficiency, tris(2-carboxyethyl)phosphine (TCEP, Thermo Scientific) was added to the purified colicin Ia to give a final concentration of 0.2 mM and it was incubated at room temperature for 30 min before adding the Alexa Fluor 488-maleimide to initiate the labelling reaction. The labelling efficiency (∼1 fluorophore per protein) was confirmed by spectrophotometry (colicin Ia: ε_280 nm_ = 59,360 M^−1^ cm^−1^; AF488: ε_495 nm_ = 71,000 M^−1^ cm^−1^) after correcting for the absorption at 280 nm by the AF488 dye (A_280 nm_ = 0.11 x A_495 nm_). The photobleaching of top-locked AF 488-labelled colicin Ia immobilised onto a quartz slide surface for TIRFM yielded a single-step drop in fluorescence intensity consistent with fluorescent-labelling at a single position.

### Production of 8-Azido-3,8-dideoxy-D-manno-octulosonic acid (Kdo-N_3_)

Kdo-N_3_ was synthesised and purified following established protocols ^18^. The synthesised Kdo-N_3_ was utilised for all ensemble and single-molecule fluorescence microscopy experiments and yielded the same results as commercially available Kdo-N_3_ (Click Chemistry Tools, Vector Laboratories, Inc.) used in more recent work. Freeze-dried stocks of the synthesised compound and the purchased compound were resuspended to a concentration of 400 mM in sterile, deionised water and stored at - 20 °C until required.

### LPS metabolic labelling with Kdo-N_3_

Pre-cultures were used to inoculate fresh medium charged with 4 mM Kdo-N_3_ to a starting theoretical OD_600_ of 0.05. Post-inoculation metabolic labelling cultures were incubated for 14 – 16 h. Cultures were subsequently harvested upon reaching late stationary phase. Cells were isolated via centrifugation (8,000 x *g*, 3 min, 4 °C) and washed three times with fresh volumes of supplemented M9 CDM containing 5 mM arabinose to remove residual Kdo-N_3_ prior to LPS fluorescent labelling.

### *In situ* fluorescent labelling of Kdo-N_3_-containing LPS inner core domains via Cu(I) catalysed azide-alkyne cycloaddition (CuAAC) in bacteria with non-capsular serotypes

Cells presenting Kdo-N_3_-containing LPS were pelleted via centrifugation and resuspended to an OD_600_ of 1.0 in ‘Click-iT’ reaction mix prepared according the ‘Click-iT’ kit protocol (‘Click-iT’ Cell Reaction Buffer Kit, Molecular Probes®, Invitrogen) supplemented with 4 mM *N*-acetylneuraminic acid (Neu5Ac) and charged with the alkyne functionalised fluorescent dye (concentrations and structures specified in Supplementary Table 8 and Supplementary Figure 11, respectively). The suspension was transferred to a sterile 2 mL microcentrifuge tube and incubated on a rotary wheel (12 rpm, 80° incline relative to bench top) for 30 min at room temperature. Post-labelling the suspension was transferred to a fresh, sterile 2 mL microcentrifuge tube and cells bearing fluorescent LPS pelleted via centrifugation. The labelled cell pellet was then washed a further three times with supplemented M9 media to remove residual fluorophore and ‘Click-iT’ mix components.

### *In vivo* fluorescent labelling of Kdo-N_3_-containing LPS inner core domains via Cu(I) catalysed azide-alkyne cycloaddition (CuAAC) in bacteria with capsular serotypes (*E. coli* UTI89 and *P. aueroginosa* PAO1)

After the three pre-labelling wash steps to remove residual Kdo-N_3_, two additional wash steps were carried out with supplemented M9 CDM containing 100 mM EDTA to remove extracellular capsular matrices and LPS for fluorescent labelling purposes. To restore native concentrations of OM co-ordinated divalent cations (Mg^2+^ and Ca^2+^) and remove residual EDTA prior to fluorescent labelling, the capsule-stripped cell pellets were then washed twice with supplemented M9 CDM. CuAAC fluorescent labelling of exposed, Kdo-N_3_-containing LPS was carried out as detailed above.

### Live cell Kdo-N_3_-containing LPS labelling of *E. coli* BW25113 or MG1655 and BW25113 Δ*waaC* via Cu(I) free strain promoted azide-alkyne cycloaddition (SPAAC)

Washed Kdo-N_3_-labelled LPS liquid culture cell samples were diluted in fresh, supplemented M9 CDM to an OD_600_ of 1.0 and transferred to a sterile 2 mL microcentrifuge tube. The appropriate volume of a dibenzocyclooctyne-amine (DBCO)-functionalised fluorophore was then added directly to this suspension (Supplementary Table 9). Labelling suspensions were then incubated for 1 h at 30 °C on a rotary wheel (12 rpm, 80° incline relative to benchtop). Suspensions were then transferred to a fresh, sterile 2 mL microcentrifuge tube and pelleted via centrifugation. The labelled cell pellet was then washed three times in freshly supplemented M9 CDM to remove residual dye. Extra care was taken during gentle resuspension of cell pellets at each stage of this protocol to ensure cells were exposed to minimal levels of shear stress thereby ensuring minimal loss of cell viability during this labelling protocol.

### *In vivo* peptidoglycan pentapeptide cross-link metabolic labelling using D-Ala-azide (D-Ala-N_3_) amino acid or propargyl-D-Ala-D-Ala (prop-D-Ala-D-Ala) dipeptide followed by fluorescent labelling via CuAAC

M9 CDM was inoculated to an OD_600_ of 0.05 via addition of an appropriate volume of MG1655 starter culture to an OD_600_ of 0.05 and charged with 4 mM D-Ala-azide or prop-D-Ala-D-Ala. Cultures were then incubated for 10 – 12 h and harvested via centrifugation upon reaching late stationary phase. The cell pellet was then washed three times to remove residual functionalised D-Ala analogue and fluorescently labelled via CuAAC via resuspension in ‘Click-iT’ mix charged appropriately functionalised AZ594-dye (Click Chemistry Tools, Vector Laboratories, Inc.; Supplementary Figure 12), selected for its ability to penetrate the outer membranes of metabolically labelled cells. ‘Click-iT’ mixes were incubated for 30 min on a rotary wheel (12 rpm, 80° incline relative to bench top) after which labelled cells were pelleted via centrifugation. Labelled cell pellets were washed three times with supplemented M9 CDM to remove residual dye and ‘Click-iT’ mix components.

### Non-canonical amino acid labelling of LptD

Sites for ncAA incorporation into LptD in *E. coli* BW25113 were selected based on *in silico* modelling of *E. coli* LptD crystal structure deposited in the PDB database [PDB: 4RHB] ^62^ (Supplementary Figure 13). Four sites for ncAA incorporation were selected based on criteria designed to minimise the impact of ncAA incorporation on LptD function and structure, whilst maximising the probability of ncAA incorporation at an accessible site to enable efficient downstream bio-orthogonal fluorophore coupling and minimise dynamic quenching of the coupled extrinsic probe by certain amino acids (*e.g*. tryptophan, histidine, methionine and tyrosine) ^63^. We selected two lysine (K473 and K602) and two aspartic acid (D592 and D600) acid residues located on two unstructured, exposed extracellular loops of LptD. These selections were informed by results from published structural, computational and biochemical studies of LptD ^64,65^. By disregarding the extracellular loops, residues and regions implicated in LptD function by these previous studies, we were able to minimise the probability that ncAA incorporation would have a deleterious impact on LptD structure and function.

Introduction of the L-lysine ncAA to the culture medium resulted in leaky expression of the recombinant, amber-stop codon-containing *lptD* gene (*lptD*)*, *pylRS* and tRNA^Pyl^ and subsequent production of recombinant, full length LptD with the ncAA integrated in one of four positions in its extracellular surface loops (LptD*). The accessible alkyne groups in the surface loops of LptD* enabled conjugation of an azide-functionalised, extrinsic fluorescent organic dye molecule via CuAAC ^20,66^. LptD* fluorescent labelling enabled *in vitro* characterisation of the recombinant protein by fluorescence imaging of LptD bands during SDS-PAGE screens. We applied the same methodology to label LptD with equivalent efficiency with ε-(tert-Butoxycarbonyl)-L-lysine (L-Lys(Boc)-OH) ncAA, (Supplementary Figure 6C) ^67,68^ when carrying out experiments that did not require LptD* fluorescent labelling. LptD is one of only two essential OMPs necessary for *E. coli* growth under standard culture conditions ^69,70^. It was therefore impossible to produce the recombinant ncAA-containing LptD proteins variants in a Δ*lptD* background. However, we were able to express recombinant ncAA-containing *lptD* variants in *E. coli* BW25113 *imp4213* ^71^. This mutant *lptD* strain produces defective LptD protein ^72^. The resulting phenotype (*e.g*. increased LPS lateral mobility, increased sensitivity to antibiotics, detergents and increased OM permeability) stems directly from the impaired function of the mutant LptD protein ^29,73^. We were able to demonstrate the functionality of the ncAA-containing LptD variants by rescuing a wild-type phenotype in the *imp4213* strain (Figure 3D and Supplementary Figure 6C), by replicating the results of previous studies in which an SDS-PAGE screen was used to show complexing of LptD with LptE (vital for LptD function and LPS insertion) (Supplementary Figure 6B) ^64,74^. Direct LptD* fluorophore conjugation via CuAAC enabled us to carry out these experiments without the use of immunoblotting for LptD and LptDE visualisation. Validation of ncAA-LptD production and subsequent fluorescent labelling were carried out for all four ncAA-containing *lptD* variants along with SDS-PAGE screens of recombinant protein production levels and ability to form a complex with LptE. Based on this screening process, the LptD^D600^ mutant was selected and used to rescue a wild-type phenotype in the *imp4213* strain (Figure 3D and Supplementary Figure 6C).

### Rescuing wild-type phenotype in the LptD-defective *E. coli* BW25113 *imp4213* strain by production of functional ncAA-containing LptD*

*E. coli* K12 BW25113 *imp4213* cells were co-transformed with pBAD-lptD* and pEVOL-pyIRS^WT^ by electroporation and selection on LB-Miller agar plates was done using reduced concentrations of ampicillin and chloramphenicol (50 μg mL^−1^ and 17.5 μg mL^−1^, respectively). Reduced antibiotic concentrations were required for selection of co-transformed cells because *imp4213* cells exhibit elevated sensitivity to antibiotics ^71,75^. A single colony of co-transformed imp4213 / pBADcLIC-lptD* / pEVOL-pyIRS^WT^ was used to inoculate a volume of LB-Miller with 50 μg mL^−1^ ampicillin and 17.5 μg mL^−1^ chloramphenicol. Post-inoculation this culture was incubated for 6 h. The pre-culture was used to inoculate two fresh volumes of LB-Miller with ampicillin and chloramphenicol antibiotics to a starting OD_600_ of 0.01. Post-inoculation cultures were incubated for approximately 3 h to an OD_600_ of 0.5. One of the cultures was charged with ε-(tert-Butoxycarbonyl)-L-lysine (L-Lys(Boc)-OH) (Fisher Scientific, Supplementary Figure 6) at a final concentration of 1 mM, and both cultures were incubated for a further 20 min. 100 μL volumes of the L-Lys(Boc)-OH supplemented culture were spread across a MacConkey agar plate and a LB/Agar with 0.1% (w/v) SDS and 5 mM EDTA plate, both supplemented with 1 mM L-Lys(Boc)-OH. Similarly, 100 μL volumes of the un-supplemented culture were spread across identical plates that had not been supplemented with L-Lys(Boc)-OH. After incubating the plates at room temperature for 5 min to allow the liquid culture to absorb into the agar, they were incubated overnight at 37 °C. Growth on L-Lys(Boc)-OH supplemented and un-supplemented plates was then compared.

### OmpA fluorescent labelling via ncAA incorporation followed by CuAAC

To investigate the distributions and spatial organisations of LPS and OMPs relative to one another in the outer leaflet of the OM, OmpA was selected for labelling via adaptation of the strategy used for LptD. OmpA is one of the most abundant OMPs in *E. coli* with approximately 100,000 molecules per cell ^4,76^, and lateral confinement of OmpA in the OM does not depend on its peptidoglycan-binding domain ^77^. A three-dimensional model of the *E. coli* OmpA structure was produced using AlphaFold2 ^78^. We then identified regions of the OmpA primary sequence predicted to form unstructured extracellular loops. We selected an accessible glutamic acid (E89) residue in extracellular loop 2 for *N*-propargyl-L-Lysine-OH incorporation via mutation of the respective codon in the *ompA* gene sequence to amber stop codons (TAG) (Supplementary Figure 5A) ^79–81^.

After isolation and amplification of the *E. coli* BW25113 *ompA* gene sequence from the purified bacterial chromosome via PCR (Supplementary Table 6), we incorporated the wild-type *ompA* gene into a pBADcLIC expression vector. The target codons in the *ompA* gene were mutated to TAG via PCR mutagenesis (Supplementary Table 6). Under standard laboratory culture conditions, OmpA is a non-essential OMP. Therefore, we were able to co-transform TAG-containing *ompA* expression vectors into an *E. coli* Δ*ompA* strain (Supplementary Table 5) along with the aaRS / suppressor tRNA vector pEVOL-pyIRS ^25^. Full length ncAA-containing OmpA (OmpA*) production was initiated by charging cell cultures with 1 mM *N*-propargyl-L-Lysine-OH. OmpA* fluorescent labelling with a functionalised small extrinsic organic dye molecule was carried out *in situ* via CuAAC.

### Dual differential fluorescent labelling of OmpA* and newly inserted Kdo-N_3_-containing LPS

*E. coli* K12 BW25113 Δ*ompA* cells were co-transformed with pBADcLIC-OmpA* and pEVOL-pyIRS via electroporation as detailed above. A 5 mL volume of supplemented M9 CDM with 100 µg mL^−1^ ampicillin, 30 µg mL^−1^ kanamycin and 35 µg mL^−1^ chloramphenicol was inoculated with a single colony of freshly co-transformed cells and incubated at 37 °C, 220 rpm agitation for approximately 7 h (to an OD_600_ of 1.5 – 2.0). This pre-culture was used to inoculate a fresh volume of supplemented M9 CDM plus selection antibiotics to a starting OD_600_ of 0.05. The culture was incubated as before for approximately 3 h, to an OD_600_ of approximately 0.5. Uninduced recombinant OmpA* production and LPS Kdo-N_3_ metabolic labelling were done simultaneously via the addition of propargyl-L-Lysine (prop-L-Lys) and Kdo-N_3_ to final concentrations of 1.0 mM and 4.0 mM, respectively. The recombinant OmpA* production / LPS metabolic labelling culture was then re-incubated under standard conditions for a further 1 h. After 1 h, the culture was removed and placed immediately on ice to halt further cell division, LPS insertion and OmpA* production. The culture was diluted with pre-chilled, supplemented M9 CDM plus selection antibiotics to an OD_600_ of 1.0 and transferred to a pre-chilled 2.0 mL microcentrifuge tube.

Cells were then harvested via centrifugation (10,000 x *g*, 3 min, 4°C) and the supernatant was carefully removed. The cell pellet was then resuspended to an OD_600_ of 1.0 in fresh, pre-chilled supplemented M9 CDM plus selection antibiotics via repeated, gentle pipetting. The resuspended culture was transferred to a fresh-prechilled 2.0 mL microcentrifuge tube on ice. This washing step was repeated three times in total to ensure removal of residual prop-L-Lys and Kdo-N_3_. The washed cell pellet was then resuspended to an OD_600_ of 1.0 in ‘Click-iT’ mix charged with AF488- or AZ647-alkyne, transferred to a 2 mL microcentrifuge tube, and incubated at room temperature for 30 min on a rotary wheel (12 rpm, 80° incline relative to benchtop). In replicate experiments, the LPS and OmpA^E89^ labelling order was alternated, as were the dye combinations (Supplementary Table 10). This alternative labelling strategy was implemented in the experimental replicates to ensure that the order in which species were labelled and the dye combinations used did not influence results (Figure 2A and Supplementary Figure 5). Cells with then pelleted via centrifugation (10,000 x *g*, 3 min, 4°C). The spent ‘Click-iT’ mix supernatant was carefully removed. The cell pellet was then resuspended to an OD_600_ of 1.0 in 0.2 µm-filtered, pre-chilled phosphate buffered saline pH 7.4 (PBS; 137 mM NaCl, 2.7 mM KCl, 4.3 mM Na_2_HPO_4_, Sigma-Aldrich) via gentle repeated pipetting.

The suspension was transferred to a fresh 2.0 mL microcentrifuge tube and cells were re-pelleted via centrifugation (10,000 x *g*, 3 min, 4 °C). The PBS supernatant was carefully removed, and the pellet resuspended to an OD_600_ of 1.0 in ‘Click-iT’ mix charged with AZ647- or AF488-N_3_. The suspension was transferred to a 2.0 mL microcentrifuge tube and incubated on a rotary wheel (12 rpm, 80° incline relative to bench top) at room temperature for 30 min. Cells were then pelleted via centrifugation (10,000 x *g*, 3 min, 4 °C). The spent ‘Click-iT’ mix was carefully removed, and the pellet resuspended in pre-chilled, filtered PBS to an OD_600_ of 1.0. The suspension was transferred to a fresh 1.5 mL pre-chilled microcentrifuge tube and cells re-pelleted via centrifugation (10,000 x *g*, 3 min, 4 °C). This washing step was repeated three times in total. Cells were then fixed via re-suspension to an OD_600_ of 1.0 in 4% (v/v) paraformaldehyde in PBS followed by incubation at room temperature for 30 min on a rotary wheel (12 rpm, 80° incline relative to benchtop). The suspension was then transferred to a fresh, pre-chilled 1.5 mL microcentrifuge tube and cells were pelleted via centrifugation (10,000 x *g*, 3 min, 4 °C). The pellet consisting of dual-labelled, fixed cells was washed three more times in pre-chilled, 0.2 µm filtered PBS as before. After the final wash step cell pellets were resuspended to an OD_600_ of 1.0 in PBS and stored at 4 °C or on ice, protected from light until imaging was done.

### LPS extraction

LPS was extracted via adaption of the hot phenol – aqueous method developed by Westphal (1965) ^82^. The cell pellet was re-suspended to a theoretical OD_600_ of 0.5 in PBS plus 0.5 mM MgCl_2_ and 0.15 mM CaCl_2_, pH 7.4 and pelleted via centrifugation (10,000 x *g*, 10 min, 4 °C). The washed cell pellets were then resuspended to the same OD_600_ in ultrapure deionised water (resistivity = 18.2 MΩ cm) and transferred to a glass dram with Teflon-coated lid. An equal volume of 90% (w/v) phenol, pre-warmed to 65 °C was added and the emulsion stirred vigorously at 65 °C for 15 min. Drams were then transferred to ice for 15 min to cool. Emulsions were then transferred to microcentrifuge tubes and centrifuged (8,500 x *g*, 10 min, 15 °C). The LPS-containing aqueous fraction was transferred to a 50 mL screw-cap polypropylene tube. A second aqueous extraction was then carried out on the remaining phenol layer and the aqueous fractions were pooled in the 50 mL screw-cap polypropylene tube. Sodium acetate was added to the pooled aqueous fractions to a final concentration of 0.5 M. 10 volumes of 95% (v/v) ethanol, pre-chilled to −20 °C was then added and after mixing via repeated inversions the suspension was incubated overnight to maximise LPS precipitation. The following day LPS was pelleted via centrifugation (2,000 x *g*, 10 min., 4 °C). After supernatant aspiration the LPS pellet was dried under a steady stream of N_2(g)_. The dried LPS pellet was resuspended in 100 µL ultrapure deionised water and transferred to a 1.5 mL microcentrifuge tube. Sodium acetate was added to a final concentration of 0.5 M and 1.1 mL pre-chilled 95% (v/v) ethanol was added. After tube content mixing via repeated inversion LPS was re-precipitated via overnight incubation at −20 °C. Precipitated LPS was pelleted via centrifugation. The pellet was dried again under N_2(g)_ and finally resuspended in 50 µL PBS.

### LPS Tricine-Sodium Dodecyl Sulfate-Polyacrylamide Gel Electrophoresis (TSDS-PAGE)

A volume (15 µL) of the purified LPS was mixed with a volume (5 µL) of 4x loading buffer (Supplementary Table 11) plus 2% (v/v) β-mercaptoethanol (β-ME). After vortexing and a brief centrifugation, the samples were incubated for 1 h at 40 °C. The samples were then re-vortexed and centrifuged briefly before loading the entire sample volume into the well of a 10 cm^2^ 1 mm thick 15% (w/v) acrylamide resolving / 6% (w/v) acrylamide stacking Tricine gel prepared according to the protocol published by Schägger (2006) ^83^. Commercial smooth LPS standards (Thermo Scientific^®^) were analysed alongside extracted LPS samples to facilitate classification of sample LPS bands. O55:B5-AZ488 LPS conjugate samples were prepared for TSDS-PAGE by adding 1 µL of 1 mg mL^−1^ LPS conjugate stock solution to 9 µL ultrapure deionised water followed by 3.3 µL of 4 x loading buffer plus 2% (v/v) β-ME (Supplementary Table 11). Samples were vortexed and briefly centrifuged before being incubated at 40 °C for 1 h, then re-vortexed and briefly centrifuged again before loading the entire sample volume into the well of a Tricine gel. Electrophoresis was carried out at 4 °C using pre-chilled buffers with the gel tank protected from light to prevent fluorophore (AF488) photobleaching. Electrophoresis was done in constant current mode for 45 min at 30 mA until a dye front formed as a fine horizontal line at the stacking / resolving gel interface. The current was then increased to 60 mA for approximately 3 h until the dye front was approximately 1 cm from the base of the gel.

### Processing and visualisation of LPS TSDS-PAGE gels

Upon completion of electrophoresis, the gels were fixed via immersion in 150 mL 50% (v/v) methanol / 3% (v/v) glacial acetic acid and incubated overnight at room temperature on a slow-moving rocker, protected from light. The following day gels were washed via immersion in 150 mL 3% (v/v) glacial acetic acid and incubated for 20 min at room temperature on a slow-moving rocker. This step was repeated three times in total. AF488-LPS bands were then visualised using an Amersham Typhoon 5 gel and blot bioimaging system equipped with a 488 nm argon laser and 525 BP filter. To visualise total LPS bands, the gels were then stained using a Pro-Q Emerald 300 LPS gel stain kit according to the manufacturer’s instructions (Thermofisher Scientific^®^). Briefly, after fixation and washing steps LPS carbohydrates were oxidised via immersion in 30 mL 1% (w/v) H_5_IO_6_ followed by incubation for 45 min on a slow-moving rocker at room temperature. Gels were then washed three times in 150 mL 3% (v/v) glacial acetic acid for 20 min. Gels were then stained via immersion in 25 mL dilute (1:50) Pro-Q Emerald 300 stain and incubation for 90 min at room temperature on a slow-moving rocker. After post-staining washing steps Pro-Q stained LPS bands were visualised under UV illumination using a GeneGenius gel imaging system.

### Pre-FRAP and pre-TIRFM treatments with chelator, detergent or urea

After *in situ* fluorescent labelling of the LPS and the post-labelling washing steps, the cell pellet was resuspended to an OD_600_ of 1.0 in M9 CDM plus chelator, *N*,*N*-Dimethyldodecylamine *N*-oxide (LDAO) or urea at pre-determined maximum concentrations cells were able to withstand whilst remaining viable and exhibiting standard growth rates (see Supplementary Table 12) and transferred to a sterile 2 mL microcentrifuge tube. The suspension was then incubated on a rotary wheel (12 rpm, 80° incline relative to benchtop) for 30 min at ambient temperature. Cells presenting deep rough and/or rough LPS (see Supplementary Table 5) were mounted and imaged in M9 CDM plus chelator, detergent or urea and 5 µm silica beads at an OD_600_ of 2.0. Cells presenting smooth LPS (see Supplementary Table 5) were mounted in PBS-charged CyGEL^TM^ (BioStatus) with 5 µm diameter silica beads added.

### Culturing bacterial cells in M9 chemically defined medium with altered Mg^2+^ and Ca^2+^ concentrations

Standard culturing, metabolic- and fluorescent labelling protocols were followed exchanging standard M9 CDM (see Supplementary Table 7) for M9 CDM supplemented with altered Mg^2+^ and Ca^2+^ concentrations (see Supplementary Table 13). Preliminary OD_600_ growth curve control experiments done in triplicate showed that exchange of standard M9 CDM for the two modified M9 compositions had no statistically significant impact on rates of cell growth.

### Mounting of *E. coli* strains producing deep rough or rough LPS for fluorescence microscopy using poly-D-Lysine coated coverslips or quartz slides

Washed and labelled cell pellets were re-suspended in freshly supplemented M9 medium (plus antibiotic(s) where required – Supplementary Table 5) to an OD_600_ of 2.0. 48 µL of the sample was mixed with 2 µL 0.5% (w/v) 5 µm diameter silica bead slurry (Bang Laboratories, slurry prepared in unsupplemented M9 CDM, final bead concentration = 0.02 % (w/v)). 30 µL of this suspension was pipetted dropwise along the long axis centreline of a clean 1.0 – 1.2 mm glass slide and covered with a 22 mm x 64 mm No. 1.5 cover slip coated with 20 μg mL^−1^ poly-D-Lysine (prepared in 50 mM MOPS, pH 8). The coverslip edges were sealed with clear nail varnish to prevent the sample drying out. The slide was then incubated in a light-excluding container for 10 min at room temperature to enable cells to adhere to the coverslip and allow the clear nail varnish to set.

### Mounting of *E. coli* strains producing smooth LPS for fluorescence microscopy using thermoreversible CyGEL

Chilled (on ice to ∼4 °C), washed and labelled smooth LPS cell pellets were resuspended to an OD_600_ of 2.0 in pre-chilled CyGEL charged with 1 x PBS pH 7.4 and 5 µm silica beads on ice via gentle, repeated pipetting. 30 µL of the suspension was then aliquoted along the long axis centreline of a re-chilled 1.0 – 1.2 mm glass slide and covered with a pre-chilled clean no. 1.5 glass coverslip. The slide was sealed via the application of a clear nail varnish bead along the edges of the coverslip and placed in a pre-chilled, light-excluding container and incubated at 4 °C for 10 min allowing even suspension dispersal within the sealed cavity between the slide and coverslip. The container was then incubated at room temperature for a further 15 min resulting in the CyGEL solidification and the satisfactory immobilisation of cells for imaging purposes.

### Fluorescence recovery after photobleaching (FRAP) and confocal fluorescence microscopy

FRAP experiments were conducted using either an upright Zeiss LSM710 confocal fluorescence microscope or an inverted Zeiss LSM 780 multiphoton confocal fluorescence microscope (in Biosciences Technology Facility, University of York) (see Supplementary Tables 14 and 15 for image acquisition settings). Data was collected using the FRAP program within the ZEN2011 software. Pre-bleach fluorescence and differential interference contrast (DIC) images were acquired. One pole was subjected to photobleaching (using a defined, standard area of 50 pixels x 30 pixels) via application of 20 - 30 laser pulses at maximum laser power. Five post-bleach images were then collected at one second intervals. Subsequent post bleach images were then collected at one-, two- and five-minute time points post-bleaching. Post-FRAP DIC images were then acquired to verify that the position of the subject cell had not changed, and the integrity of its cell envelope had not been compromised.

### Quantitative analysis of FRAP data

Data from individual FRAP image sequences were converted into recovery curves by plotting normalised fluorescence intensity in the bleached region over time. Normalisation of fluorescence recovery was necessary for comparison of individual FRAP analyses since levels of initial fluorescence, and the extent of photobleaching varied from cell to cell. A double-normalisation technique was then applied to normalise the fluorescence intensity in the bleached region of each image against both the initial cell fluorescence and the degree of sample photobleaching during image acquisitions ^84^.

#### For each time series image

Bleached region average fluorescence intensity (***B_t_***) and whole cell average fluorescence intensity (***T_t_***) values are normalised against corresponding background average fluorescence intensity (***BG_t_***):

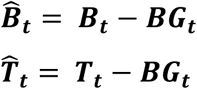

At each time point, ***t*** the double normalised average fluorescence intensity in the bleached region (***I***^_***t***_) is obtained using the formula below:

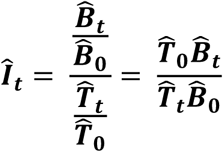

Recovery curves were constructed from FRAP data using a custom MATLAB code (available at https://github.com/RosieLeaman/FRAP). The FRAP image sequence data was processed in the following steps:

1. Identification of the cell being photobleached (if multiple cells are in the image).
2. Identification of the bleached area boundaries, cell outline and identification of a region of background that does not contain any other cells for the purposes of double normalisation.
3. Measurement of the average intensity in each of the three regions at each time point, applying any drift correction in the *xy* plane as required.
4. Construction of the corresponding recovery curve from the extracted average intensities at each time point (see Supplementary Figure 14).

Mobile fraction formula:

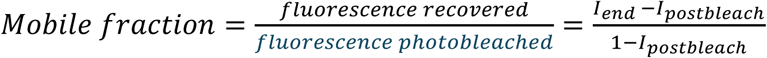

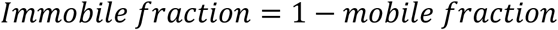

*I* = average fluorescence intensity in bleached region.
*I_end_* = average fluorescence intensity in bleached region at 300 s.
*I_postbleach_* = average fluorescence intensity in bleached immediately after photobleaching step.

### Statistical analysis of FRAP data

The mobile fraction was defined as the proportion of labelled species (in this context AZDye-labelled, Kdo-analogue containing LPS) that undergo lateral diffusion over the course of the experiment. The immobile fraction was defined as proportion of molecules that do not undergo diffusion over the course of the experiment. As detailed in Supplementary Figure 14, the mobile fraction of fluorescently labelled LPS molecules can be derived from the FRAP recovery curves. Prior double normalisation enabled comparison between different experimental conditions. Individual experiment mobile fractions were calculated in Microsoft Excel using the following equation and data outputs from the FRAP analysis code implemented in MATLAB (Mathworks):

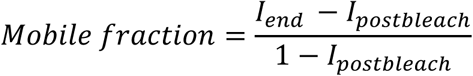

For each condition, the labelled LPS mobile fractions was calculated in at least 30 individual cells (n ≥ 30) collected over at least 3 experimental replicates (N ≥ 3). Box plots were produced representing the mobile fraction distributions for given conditions using the seaborn Python library. The box plots showed the median mobile fraction and interquartile range with whiskers extending to the maximum and minimum values. Values outside of the whiskers were determined as outliers by the seaborn box plot function. To assess differences in the mobile fraction distributions obtained from two conditions a Mann-Whitney U-test was applied since it does not assume that the underlying distribution is normal. The test was applied in MATLAB using the ranksum function. Differences in the mobile fraction distributions of two conditions were judged statistically insignificant if the *p* value of the test > 0.05.

### Preparation of labelled cell samples for direct stochastic optical reconstruction microscopy (dSTORM)

Immediately after the ‘Click-iT’ labelling step, the cell suspension was transferred to a pre-chilled 2 mL microcentrifuge tube and pelleted via centrifugation (10,000 – 12,000 x *g*, 2 min, 4 °C). Labelled cell pellets were re-suspended to an OD_600_ of 1.0 in 0.2 µm filtered PBS supplemented with 2 mM MgSO_4_ and 0.5 mM CaCl_2_, and transferred to a fresh, pre-chilled 2 mL microcentrifuge tube. Cells were re-pelleted via centrifugation. This washing step was carried out a total of three times. The washed and labelled cell pellet was fixed via resuspension to an OD_600_ of 1.5 in 4% (v/v) ultrapure, methanol-free paraformaldehyde (PFA, Polysciences Ltd.) in PBS. This solution was incubated for 30 min on a rotary wheel (12 rpm, 80° incline relative to benchtop) at room temperature. The fixed cell suspension was then transferred to a fresh 1.5 mL microcentrifuge tube and pelleted via centrifugation. The pellet of fixed and labelled cells was resuspended to an OD_600_ of 1.0 in PBS with MgSO_4_ and CaCl_2_, transferred to a fresh microcentrifuge tube and pelleted via centrifugation. This washing step was carried out three times in total.

### Mounting of fixed, washed and fluorescently labelled cell samples using poly-D-Lysine coated coverslips for imaging by dSTORM

The washed, fixed, and labelled cell pellet in a microcentrifuge tube was placed on ice to cool to 4 °C and resuspended in degassed, pre-chilled Glucose oxidase (GluOx) (Sigma Aldrich) / Catalase (Cat) (Sigma Aldrich) O_2_ scavenging buffer to an OD_600_ of 1.5 via repeated gentle pipetting and vortexing (Supplementary Table 16) ^85,86^. 45.5 µL of the suspension was transferred to a 500 µL microcentrifuge tube on ice. 2.5 µL of 1 M β-mercaptoethylamine hydrochloride (β-MEA) (50 mM final concentration), 2 µL of 0.5% (w/v) 5 µm diameter silica bead slurry (0.02% (w/v) final bead concentration) and 0.5 µL TetraSpek 0.2 µm microsphere standard solution (1.5 × 10^9^ particles mL^−1^, ThermoScientific) were added. TetraSpek beads were added to enable accurate AF488 and AZ647 channel alignment during two-colour dSTORM image processing. 10 µL of the suspension was then pipetted onto the centre of a pre-chilled (to minimise O_2_ scavenging buffer enzyme activity), clean 1.0 – 1.2 mm thick glass slide. The slide was covered with a poly-D-lysine-coated, 18 mm^2^ high precision (Zeiss), no. 1.5 glass coverslip ensuring no bubbles were present in the slide chamber. The chamber was sealed with a clear nail varnish. The slide was then incubated at 4 °C for 20 min while protected from light to allow the nail varnish to dry and the cells to adhere to the poly-D-lysine-coated coverslip surface.

### 2D dSTORM data acquisition with AF488- and AZ647-labelled samples

dSTORM imaging experiments was done on a Zeiss Elyra 7 super-resolution imaging platform in laser widefield beam path mode using ZEN Black 3.0 SR FP2 software (Biosciences Technology Facility, University of York). Objective coupled total internal reflection fluorescence (TIRF) illumination was employed using a Plan-Apochromat 63 x / 1.46 NA Korr oil immersion objective Var 2 lens together with TIRF uHP (ultra-high power) laser power density setting. Image areas were set to 128 pixels x 128 pixels for two cells or 64 pixels x 64 pixels for a single cell and saved in 16-bit format. Initial excitation and intersystem crossing of all fluorophores in the target cell(s) from their ground singlet state to an excited triplet ‘dark’ state was achieved via application of a short (200 – 300 frames, 50 ms exposure time), high intensity laser pulse (488 nm: 20-25%, 642 nm: 10-15%) with the microscope in epifluorescence (EPI) mode. dSTORM time series image sequences were then collected using highly inclined and laminated optical sheet (HILO) illumination for 10,000 frames (50 ms exposure time) using reduced laser powers (488 nm: Starting at 2% increasing to 15% over the time series, 642 nm: Starting at 0.7% and increasing to 5% over the time series). To propagate fluorophore ‘blinking’ in the latter 5000 frames, 405 nm light was introduced during the ‘transfer’ phase of individual image collection to reduce fluorophore photobleaching resulting from simultaneous exposure to both 488 nm and 405 nm light ^86^. HILO illumination mode was used to enable excitation of fluorophores in the outer membrane furthest from the coverslip thereby maximising the signal to noise (S:N) ratio. Additional dSTORM image acquisition settings are summarised in Supplementary Table 17.

Two-colour dSTORM imaging of AF488- and AZ647-labelled species in the OM of in-frame, in-focus cells were done sequentially using the Zeiss Elyra 7 super-resolution imaging platform with 488 nm and 642 nm excitation lasers. The BP420-480 + BP490-550 emission filter (Supplementary Table 17) was used to prevent cross channel ‘bleeding’ of fluorescence emission from the two dyes. dSTORM data for AZ647-labelled species was collected first using the second sCMOS camera due to the lower photostability of the dye compared to AF488. dSTORM data for AF88-labelled species was then collected using the first sCMOS camera. After the initial high intensity burst to trigger fluorophore intersystem crossing from ground singlet states to a ‘dark’ triplet state, the fluorescent emission ‘blinking’ events were collected over 10,000 frames using a 50 ms exposure for each channel.

### Processing of single colour channel 2D-dSTORM images

dSTORM raw data were processed using the SMLM (single-molecule localisation microscopy) processing facility found within the ZEN Black 3.0 SR software. Time series image sequences were converted to a crude single dSTORM image discarding overlapping molecules using a x,y Gauss fit model, with a peak mask size of 9 pixels and a peak intensity to noise ratio of 6.5. Typical filtering settings are detailed in Supplementary Table 18. SMLM-grouping settings were adjusted to minimise double counting of individual dye molecules (5 frames for the maximum “ON” time, 50 frames for maximum ‘OFF’ gap, 1.7 pixel for capture radius). Pixel resolution was set at 10 nm pixel^−1^ with localised peaks displayed in gauss mode reflecting the degree of localisation precision. dSTORM image data was converted into images in .CZI format using the ZEN 3.0 SR software ‘Convert to Image’ tool and exported to FIJI for final image production ^87^. Final images were saved in both .TIFF and .PNG formats.

### Co-localisation analysis of fluorescently labelled newly inserted LPS and propargyl-OmpA^E89^ in the bacterial OM

Initial two-colour dSTORM image processing and subsequent production was carried out for each colour channel in ZEN black Elyra software as detailed above. Channels were aligned using the ‘Channel Alignment’ tool in ZEN black 3.0 SR software using the positions of TetraSpek microspheres are markers. TetraSpek microspheres also served as fiducials for drift correction during SMLM image processing. Subsequent image processing and co-localisation analyses were carried out in FIJI ^87^. Each two-colour 2D dSTORM cell image was processed individually using a standard co-localisation method. A Gaussian blur was applied to both channels (σ = 1.0). The channels were separated into individual images and converted to TIFF format (8 bit) and converted to a greyscale colour scheme.

Regions for co-localisation analysis were selected via application of a minimal threshold based on the image with the lowest apparent signal (usually fluorescently labelled propargyl-OmpA^E89^) to exclude extracellular regions and dark regions, thus preventing background fluorescence signal and its ‘absence’ from artificially inflating correlation values. The Co-loc2 FIJI plug-In was then applied to obtain Pearson correlation coefficient (PCC) values describing the degree of co-localisation of the labelled species in the 2 channels ^88,89^. This method for co-localisation analysis was selected because PCC is independent of signal and background fluorescence levels removing the need for extended image pre-processing and thus minimising the potential influence from unconscious user bias. Normalised output values (+1 to −1) enable results for individual cells to be grouped and PCC value distributions from different conditions compared without further processing. The Co-loc2 plug-In superimposed the fluorescent distributions of both channels represented by scatterplots. The application of linear fits to these scatterplots enabled the calculation of PCC values for each two-colour image describing the pixel-by-pixel covariance in both channels.

The PCC formula is given by:

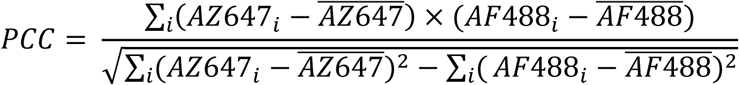

*AZ647i* = AZ647 fluorescent intensity in pixel *i*
*AF488i* = AZ488 fluorescent intensity in pixel *i*
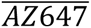 = AZ647 mean fluorescent intensity
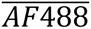 = AZ488 mean fluorescent intensity

Potential PCC values range from +1 where the fluorescent intensities of the two channels are perfectly linearly related (*i.e*. perfect co-localisation) to −1 where fluorescent intensities are perfectly inversely related (*i.e*. a complete absence of co-localisation). A value of 0 is expected if the distributions of the two differentially labelled fluorescent species were independent of one another.

### Identification and measuring of LPS and OmpA containing regions in 2D two-colour dSTORM images

After processing of the two-colour 2D-dSTORM data and conversion to .CZI images, the LPS and OmpA channels were split into separate images in ImageJ (version 1.54j) ^90^. A global threshold was applied to each image using the ISODATA auto threshold algorithm to define discrete patches within each image. The image was then converted to a binary mask using the ‘Make binary’ tool in ImageJ. Lateral and perpendicular longitudinal measurement was recorded for patches with median diameters > 50 µm. Histograms of the patch dimensions were prepared using Origin (OriginLab).

### *In vivo* TIRFM imaging and single-particle tracking of fluorescently labelled LPS or colicin Ia/CirA complex

For localisation of LPS, the bacteria were metabolically labelled with Kdo-azide and fluorescently labelled using AFDye 488 DBCO copper-free SPAAC as described above (Supplementary Table 9). For localisation of CirA receptors, the bacterial cells were resuspended in CDM (OD_600_ ≈ 1) containing the AF488-labelled colicin Ia probe at a concentration of 300 nM. The suspension was then incubated on a rotary wheel (12 rpm, 80° incline relative to benchtop) for 30 min at room temperature. The labelled cells were washed twice by centrifugation (10,000 – 12,000 x *g*, 2 min) in a 2 mL microcentrifuge tube and re-suspended in a reduced volume of fresh supplemented M9 CDM to yield a final theoretical OD_600_ of 3-4. Bacteria were immobilised on poly-D-lysine coated quartz sides in supplemented M9 CDM for TIRFM using an ultra-thin sample chamber as described previously ^33^.

The prism-coupled TIRFM (including the 488 nm laser power, optical filters and two-channel image splitter) used in this work was described previously ^33^. All digital video was recorded with an Evolve 512 emCCD (Photometrics) at 30 Hz (512 x 512 pixels^2^) or at 67 Hz (128 x 128 pixels^2^) and collected at room temperature (20-22 °C). Video data was collected after moderate photobleaching to enable the detection and tracking of single fluorophores. All raw digital video was processed with a moving 3-frame median time filter implemented in MATLAB (MathWorks), then analysed with PaTrack software to localise (with Gaussian fitting of fluorescence peak to achieve sub-pixel resolution) and track fluorophores ^91^. This tracking software uses a back-propagation neural network trained on synthetic trajectories to assign free Brownian, confined and directed diffusion modes in each single-particle trajectory. The algorithm was particularly useful in this work where individual LPS molecules might display both free Brownian and confined diffusion characteristics in single trajectory. The single-molecule trajectories analysed in this work were all at least 0.9 s in duration (typically 0.9 – 2 s in duration) and displayed either confined or free Brownian diffusion (*i.e*. mixed trajectories were not observed). The type of diffusion assigned to the trajectories was confirmed by analysing the categorised trajectories again using TrackMate ^92^. The following parameters were routinely used in PaTrack for tracking fluorophores in the video data (at 30 Hz and 67 Hz) for AZDye 488-labelled rough or deep rough LPS and AF488-labelled colicin Ia/CirA receptor complex: resolution = 0.096 µm/pixel, dimension particle = 0.4 µm, maximum particle displacement = 0.4 µm, death frames = 5, short trajectory filter = 12 frames, and maximum eccentricity = 1.7. The fluorescence intensity of every tracked fluorophore was manually checked to confirm that photobleaching occurred in a single-step drop to the background intensity level which indicated a single fluorophore was present.

Fluorophores with a multi-step drop in fluorescence intensity during photobleaching and those that did not photobleach during the video recording (thus precluding use of the aforementioned test) were omitted from the MSD analysis. Once tracked, any fluorophores localised to a bacterial cell with an end-of-trajectory, asymptotic MSD value greater than 0.008 µm^2^ were retained (MSD_end of trajectory_ > 0.008 µm^2^), while those with a lower end-of-trajectory MSD value (MSD_end of trajectory_ < 0.008 µm^2^) were considered immobilised on the quartz surface and discarded. The confinement diameter (*d*_confine_) was calculated for each track by averaging MSD data for a specific time window (time lag ≈ 0.5-0.8 s) and inputting this average MSD (MSD_average_) into the following equation ^93^: MSD_average_ = (d_confine_/2)^2^/6. The two-dimensional diffusion coefficient (D_2D_) was calculated from the time-dependence of the MSD using linear regression of the first 4 time delays for confined particles or the first 13 time delays for freely diffusing particles.

## Acknowledgements

We thank the University of York Bioscience Technology Facility (BTF) for access to microscopy facilities, and G. Calder (BTF) and N. Sergent (Zeiss) for expert assistance with dSTORM data collection and analysis. We thank the following researchers for providing bacterial strains or plasmid DNA constructs: D. Browning and I. Henderson (OAR *E. coli* DFB1655), M. van der Woude (*S*. Typhimurium LT2 and *E. coli* UTI89), G. Thomas (*P. aeruginosa* PA01), E. Severi (*E. coli* BW25113 with *imp4213* mutation), P. de Boer (pNP4 plasmid) and E. Lemke (pEVOL-pylRS plasmid). We thank C. Sharrock and B. Tello Rubio for assistance with site-directed mutagenesis in LptD and OmpA, respectively. We thank P. Fogg, C. Hill, C. Spicer and G. Thomas for comments on the manuscript. C.G.B. thanks C. Kleanthous and S. Khalid (University of Oxford) for helpful research discussions and sharing unpublished data. This work was supported by The University of York, the BBSRC (19ALERT Mid-Range Equipment Initiative Award to the Department of Biology to purchase ZEISS Elyra 7 system, BB/T017589/1), the BBSRC White Rose Doctoral Training Partnership (PhD studentship award to J.N., 2272649), the Wellcome Trust CIDCATS Interdisciplinary PhD Training Programme (PhD studentship award to R.M.L., WT095024MA), a Horizon Europe Guarantee Award to M.A.F. (selected by the ERC and funded by UKRI, EP/X023680/1) and a MRC Discovery Award to C.G.B. and M.C.C. (MC-PC-15073).

## Author contributions

J.N., R.M.L., S.L., M.A.F. and C.G.B. designed the experiments with input from D.O.P. and M.C.C. J.N. and R.M.L. collected and analysed the FRAP data with assistance from C.G.B. R.M.L. and S.L. collected and analysed the SPT-TIRFM data with assistance from C.G.B. J.N. characterised the fluorescently labelled LPS glycoforms and OMPs. R.J.S. and M.A.F. synthesised and purified the Kdo-azide. L.M. and C.G.B. purified colicin Ia and labelled with fluorophore. J.N. collected and analysed all dSTORM data with assistance from C.G.B. J.N. prepared all figures with input from M.A.F. and C.G.B. J.N., M.A.F. and C.G.B. wrote the manuscript. All co-authors had the opportunity to comment on the final submitted version of the manuscript.

## Supplementary Information

**Supplementary Figure 1:**
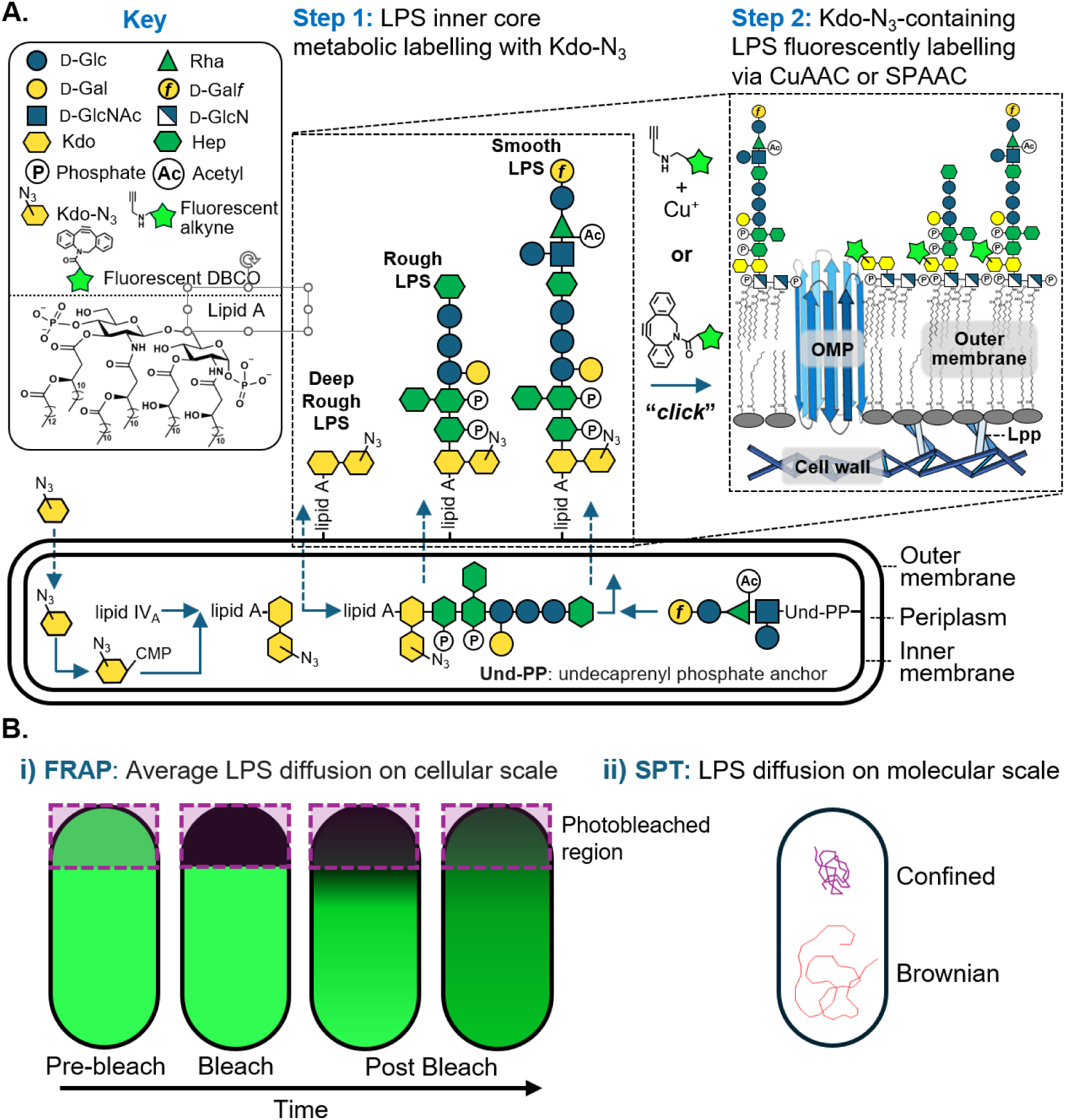
Efficient and specific fluorescent-labelling of LPS via a 2-step bio-orthogonal approach enabled average LPS lateral mobility to be assessed via *in vivo* fluorescence microscopy. **A.** Fluorescent labelling of LPS at the cell surface. *In situ* metabolic labelling of the chemically conserved LPS inner core oligosaccharide was done with an azide functionalised Kdo-analogue. Azide handles within LPS inner core oligosaccharide domain enabled bio-orthogonal conjugation of alkyne-functionalised small, photostable organic fluorescence dyes via Cu(I)-catalysed (CuAAC) or Cu(I)-free strain promoted azide-alkyne cycloaddition (SPAAC). **Gal*f*:** Galactofuranose, **Glc:** Glucose, **Rha:** Rhamnose, **GlcNAc:** *N*-Acetyl-glucosamine, **Hep:** Heptose, **Kdo:** 3-Deoxy-D-manno-oct-2-ulosonic acid, **Gal:** Galactose, **GlcN:** Glucosamine, **IM:** Inner membrane, **Lpp:** Braun’s lipoprotein, **OMP:** Outer membrane protein, **Und-PP:** undecaprenyl pyrophosphate anchor. **B.** Characterisation of LPS lateral mobility by fluorescence microscopy. LPS lateral mobility was assessed in a range of Gram-negative bacterial strains under different conditions using (i) fluorescence recovery after photobleaching (FRAP) and (ii) single-particle tracking (SPT) after detection by total internal reflection fluorescence microscopy. The photobleached region of the cell in the schematic for the FRAP experiment has a dashed outline and violet shading. Both confined and free Brownian lateral diffusion could be detected in the SPT experiments, and the type of lateral diffusion observed was found to vary depending on the strain and treatment type.

**Supplementary Figure 2:**
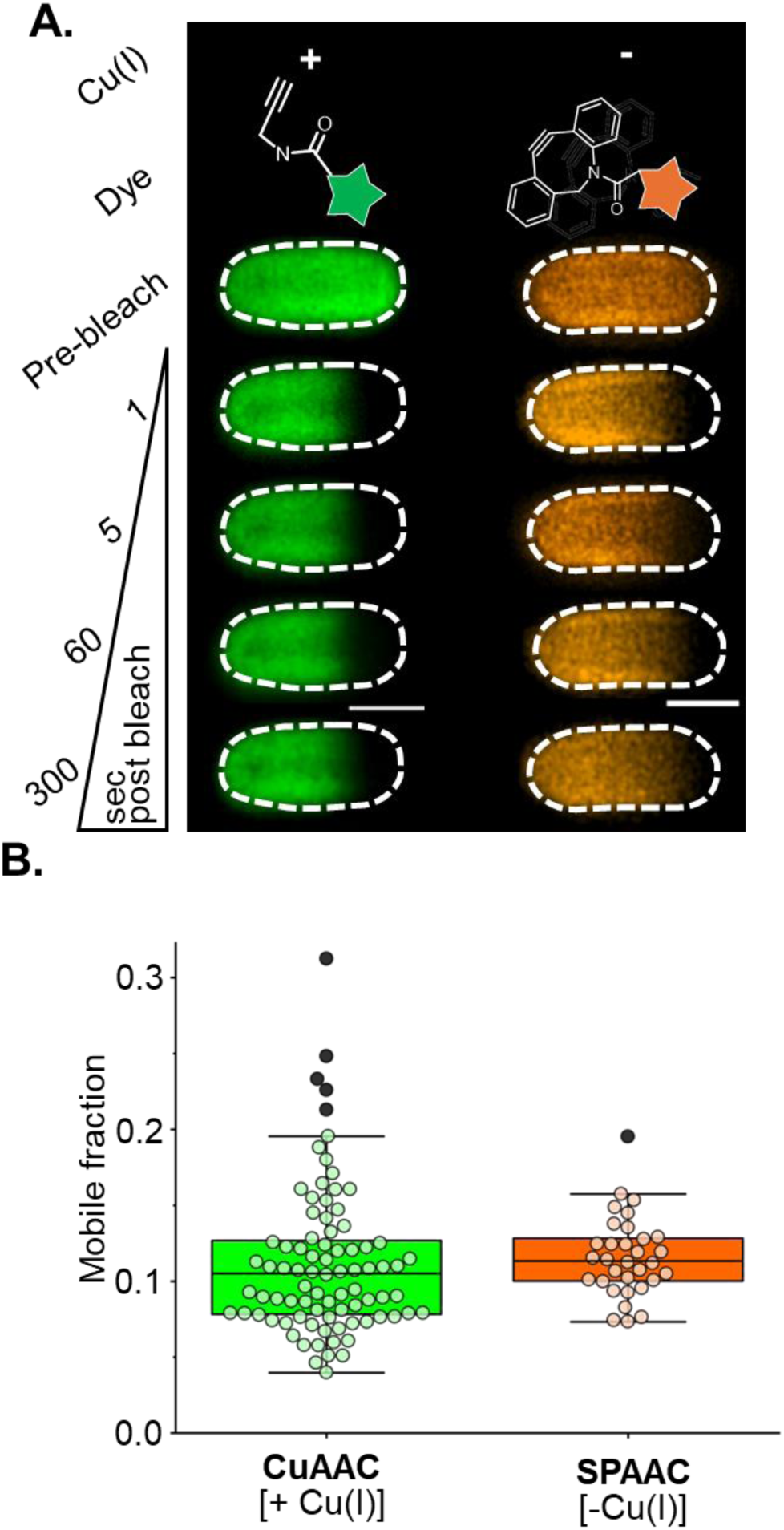
Exposure to Cu(I) during fluorescent labelling of LPS does not significantly affect LPS lateral mobility as measured by FRAP. **A.** Representative FRAP time lapse images monitoring fluorescence recovery in photobleached regions of the OM for Δ*waaC* cells with LPS labelled using AF488-alkyne via CuAAC (left) or AF568-DBCO via SPAAC (right). Scale bars: 1.0 µm. **B.** No statistically significant difference (by Mann-Whitney test, *p* = 0.11) was observed in the LPS mobile fractions in the OM of *E. coli* Δ*waaC* cells irrespective of whether LPS was labelled via Cu(I)-catalysed azide-alkyne cycloaddition (CuAAC, green box) or Cu(I)-free strain promoted azide-alkyne cycloaddition (SPAAC, orange box). CuAAC: median mobile fraction = 0.11 (n = 84). SPAAC: median mobile fraction = 0.11 (n = 32). Whiskers represent maximum and minimum values, and each coloured dot represents an individual measurement collected from at least three experimental replicates. Dark grey dots outside whisker bounds are classed as outliers.

**Supplementary Figure 3:**
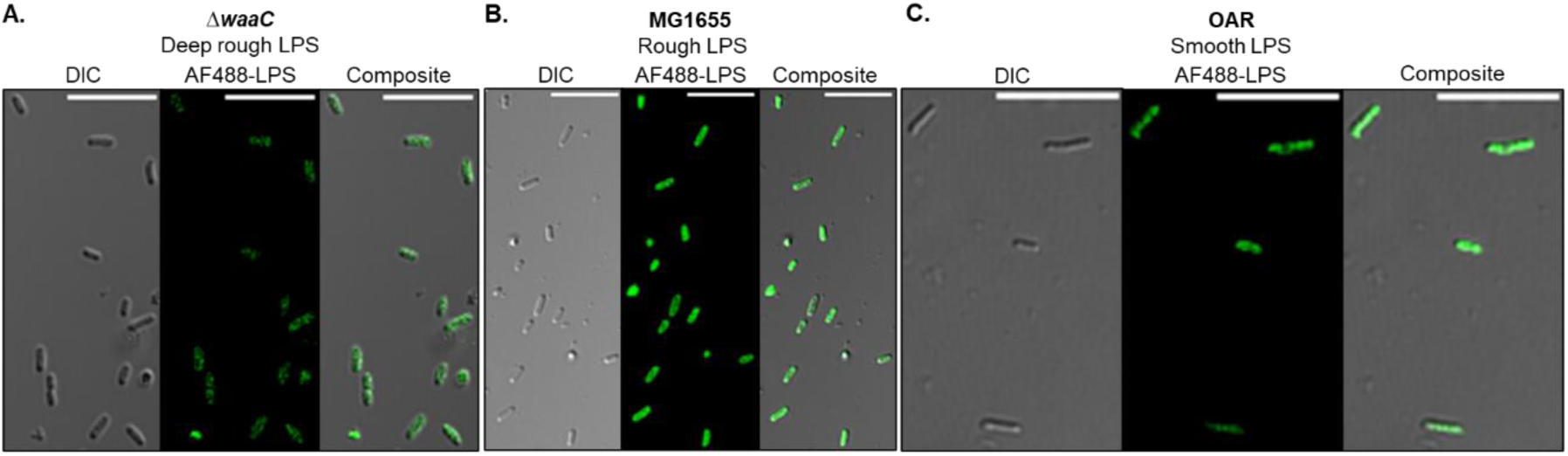
Wide-field confocal fluorescence microscopy images showing efficient *in vivo* fluorescent labelling of bacterial cells with different LPS glycoforms. **A.** *E. coli* Δ*waaC* cells producing deep rough LPS, **B.** *E. coli* MG1655 cells producing rough LPS and **C.** *E. coli* DFB1655 cells producing smooth LPS (OAR). The different LPS glycoforms were labelled using a two-step metabolic / bio-orthogonal labelling approach involving i) metabolic labelling of LPS using an azide functionalised Kdo-analogue followed by ii) *in situ* AF488-alkyne conjugation via Cu(I)-catalysed azide-alkyne cycloaddition. Scale bars: 10 µm.

**Supplementary Figure 4:**
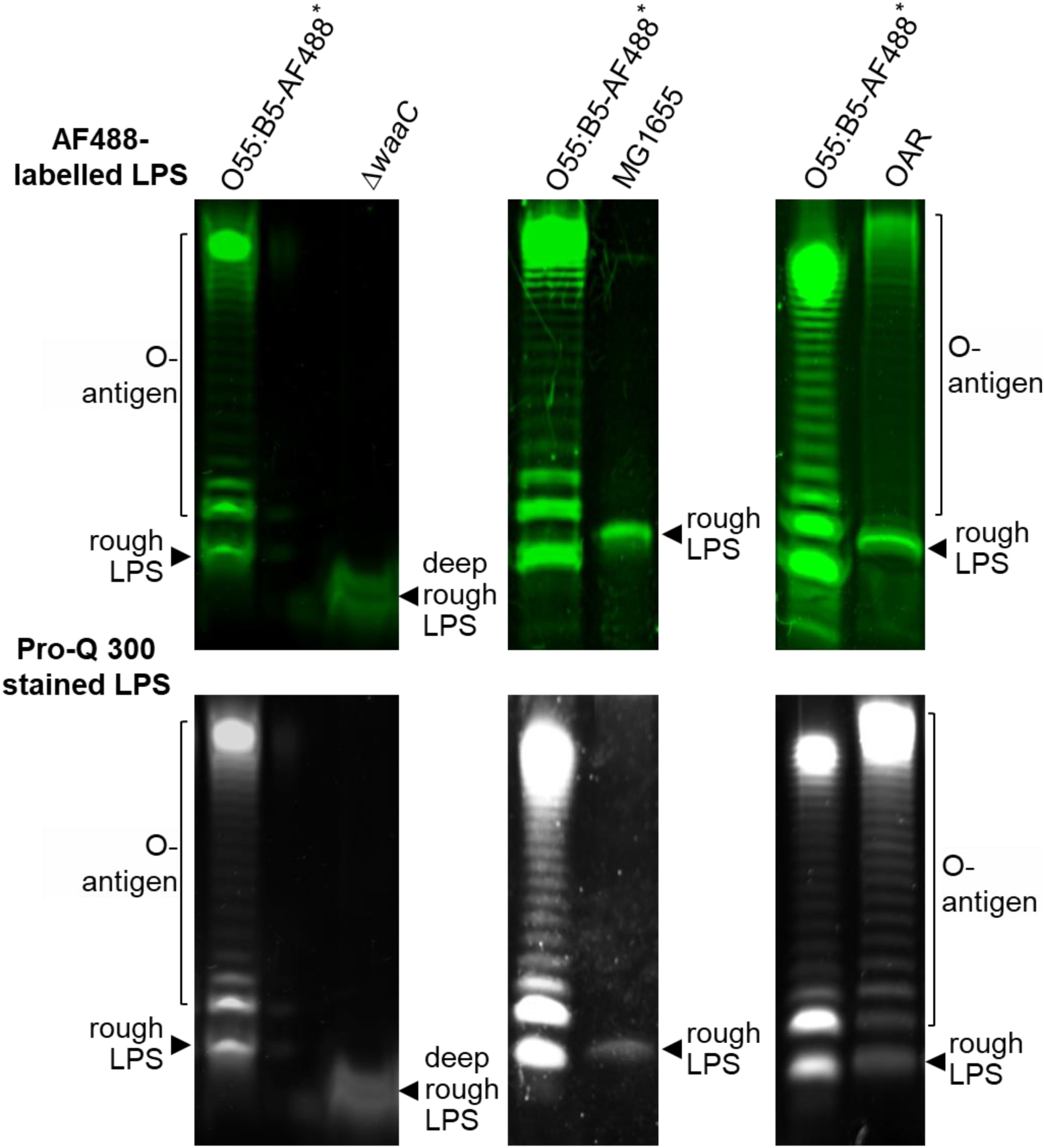
TSDS-PAGE analysis of LPS extracted from bacterial cells after two-step metabolic / bio-orthogonal labelling with Kdo-N_3_ and AF488-alkyne. LPS was extracted from *E. coli* Δ*waaC* (left-hand gel images), *E. coli* MG1655 (middle gel images) and *E. coli* DFB1655 O-antigen restored (OAR) cells (right-hand gel images). AF488-labelled LPS was visualised with 488 nm laser excitation (top row of gel images) and total LPS was visualised using Pro-Q Emerald 300 LPS staining with UV illumination (bottom row of gel images). *Purchased AF488-labelled O55:B5 LPS standard (0.1 µg per lane).

**Supplementary Figure 5:**
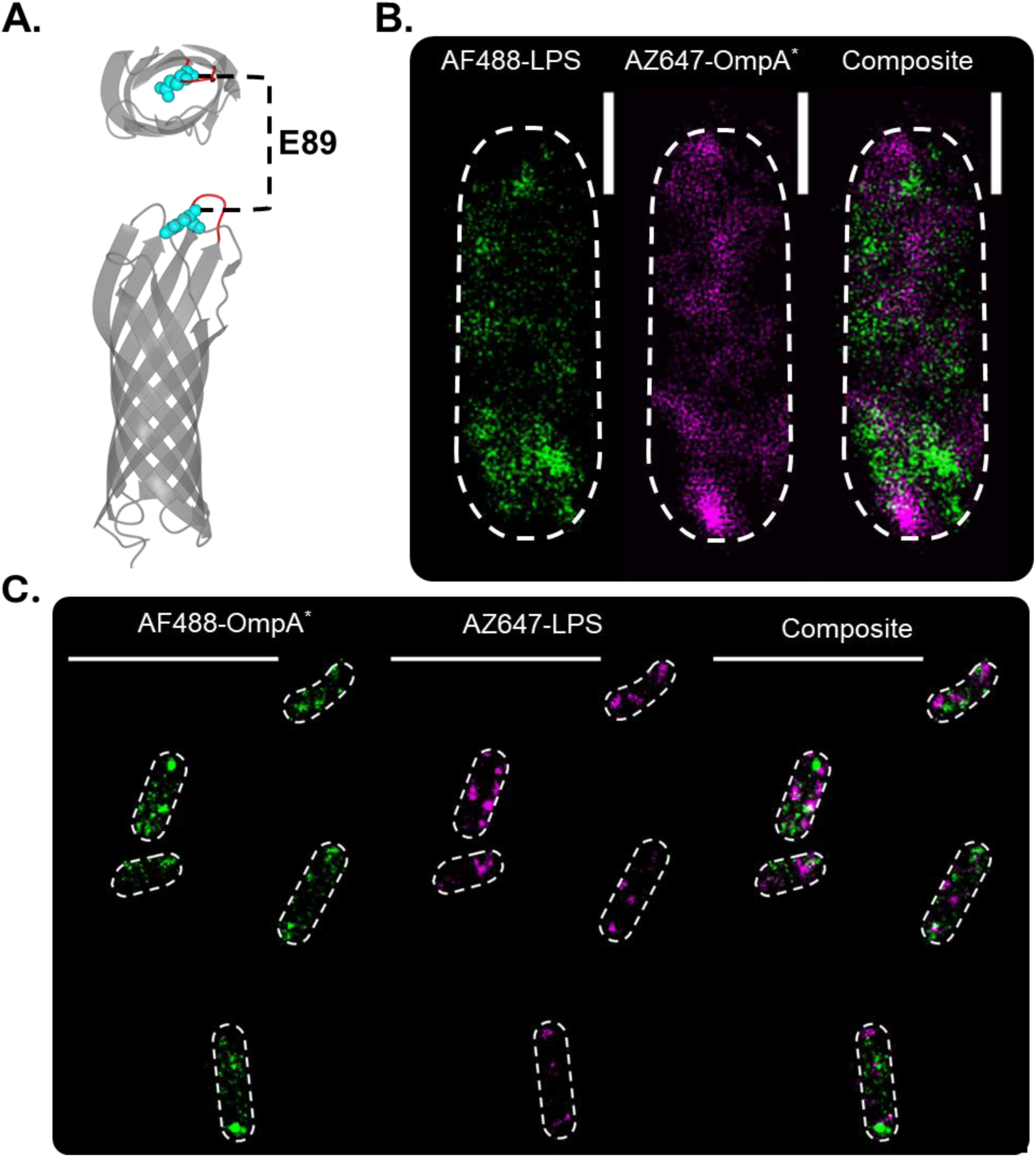
*In situ* fluorescent labelling of OmpA using amber stop codon suppression and genetic code expansion followed by fluorophore conjugation via CuAAC enabled simultaneous visualisation of discrete OMP and LPS distributions in the OM of single cells by two-colour dSTORM. **A.** AlphaFold2 generated 3D structure of *E. coli* OmpA highlighting the site of *N-*propargyl-L-lysine incorporation which replaces a glutamic acid residue (E89) in extracellular loop 2. **B.** Two-colour single-cell dSTORM images of *E. coli* Δ*ompA* cells producing recombinant OmpA* and incorporating Kdo-azide modified LPS. In the single-cell view, the separate images show the discrete positions of AF488-labelled LPS molecules and AZ647-labelled OmpA* proteins in the OM, and the composite image shows that the spatial distribution of these two OM components is heterogeneous. **C.** Two-colour wide-field dSTORM images of multiple *E. coli* Δ*ompA* cells producing recombinant OmpA* and incorporating Kdo-azide modified LPS after dual fluorescent labelling. The images of AF488-OmpA* and AZ647-LPS in the OM show that the spatial distribution of the LPS- and OMP-rich regions varies between cells, and does not depend on the fluorophore used to localise LPS and OmpA*. Scale bars in wide-field view = 5.0 µm. Scale bars in single-cell view = 0.5 µm.

**Supplementary Figure 6:**
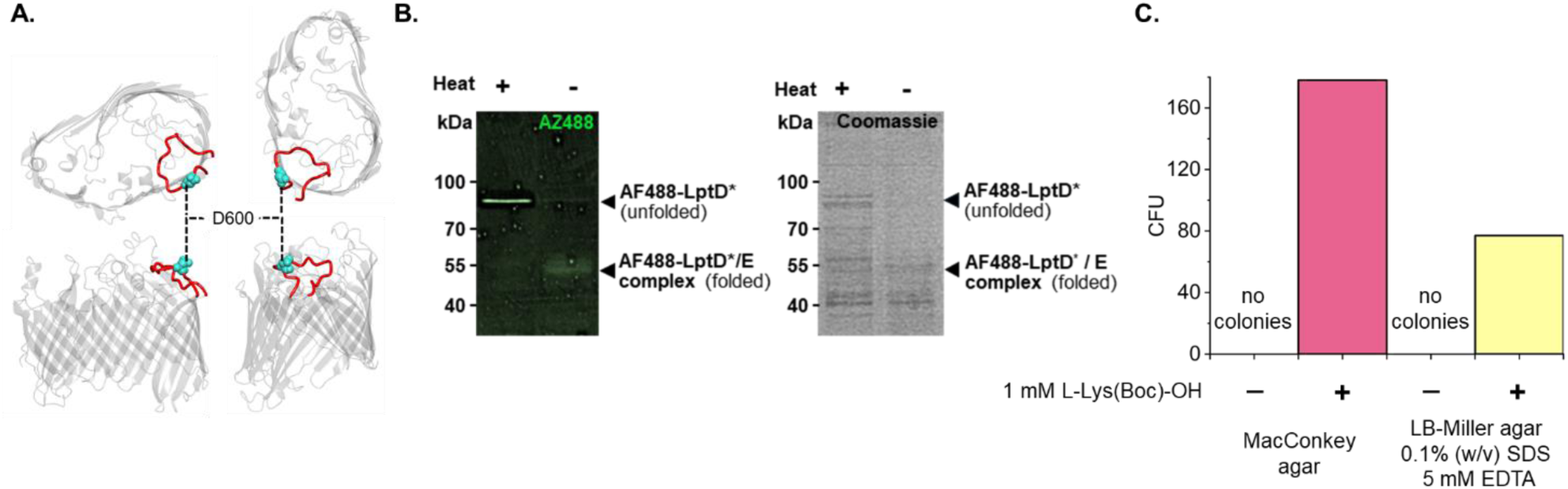
Incorporation of ε-(tert-Butoxycarbonyl)-_L_-lysine or *N*-propargyl-_L_-lysine into extracellular loop 9 of *E. coli* LptD via amber stop codon suppression and genetic code expansion. **A.** LptD crystal structure [PDB: 4HRB] highlighting loop 9 (red) and position of D600 where the site of amber stop codon mutation was introduced and the ncAA was incorporated. **B.** Production of functional LptD was regulated by addition of the following inert ncAAs: ε-(tert-Butoxycarbonyl)-_L_-lysine (_L_-Lys(Boc)-OH) allowed biochemical control over full-length LptD translation, and clickable _L_-Lys(Propargyl)-OH enabled subsequent fluorescent visualisation of protein bands for LptD* and the native LptD*/LptE complex by SDS-PAGE. SDS-PAGE was used to characterise cell lysates containing folded (unheated, lanes with ‘minus’ sign) or denatured (heat treated, lanes with ‘plus’ sign) fluorescently labelled recombinant LptD*. *N*-propargyl-_L_-lysine incorporation into LptD* followed by *in situ* fluorescent labelling with AF488-alkyne via CuAAC does not disrupt native complex formation with LptE demonstrating ncAA-LptD* is functional. Heat denaturation of the cell lysate disrupted the LptD* / LptE complex and resulted in the reduced mobility of denatured LptD* during SDS-PAGE. The SDS-PAGE gel was imaged with 488 nm laser excitation to view fluorescently labelled LptD* band (left), and with visible light after Coomassie brilliant blue staining to view all protein bands present (right). **C.** Production of _L_-Lys(Boc)-OH containing LptD* in *imp4213* cells co-transformed with pBADcLIC-lptD* and pEVOL-pylRS plasmids restored wild-type LPS insertion rates in the OM, OM asymmetry and OM barrier function. This made *imp4213* cells co-transformed with pBADcLIC-lptD* and pEVOL-pylRS plasmids tolerant to normally lethal concentrations of bile salts (MacConkey agar) and detergents (SDS) / chelators (EDTA) when grown with 1 mM _L_-Lys(Boc)-OH. Colony forming units (CFU) for a representative growth screening experiments are provided.

**Supplementary Figure 7:**
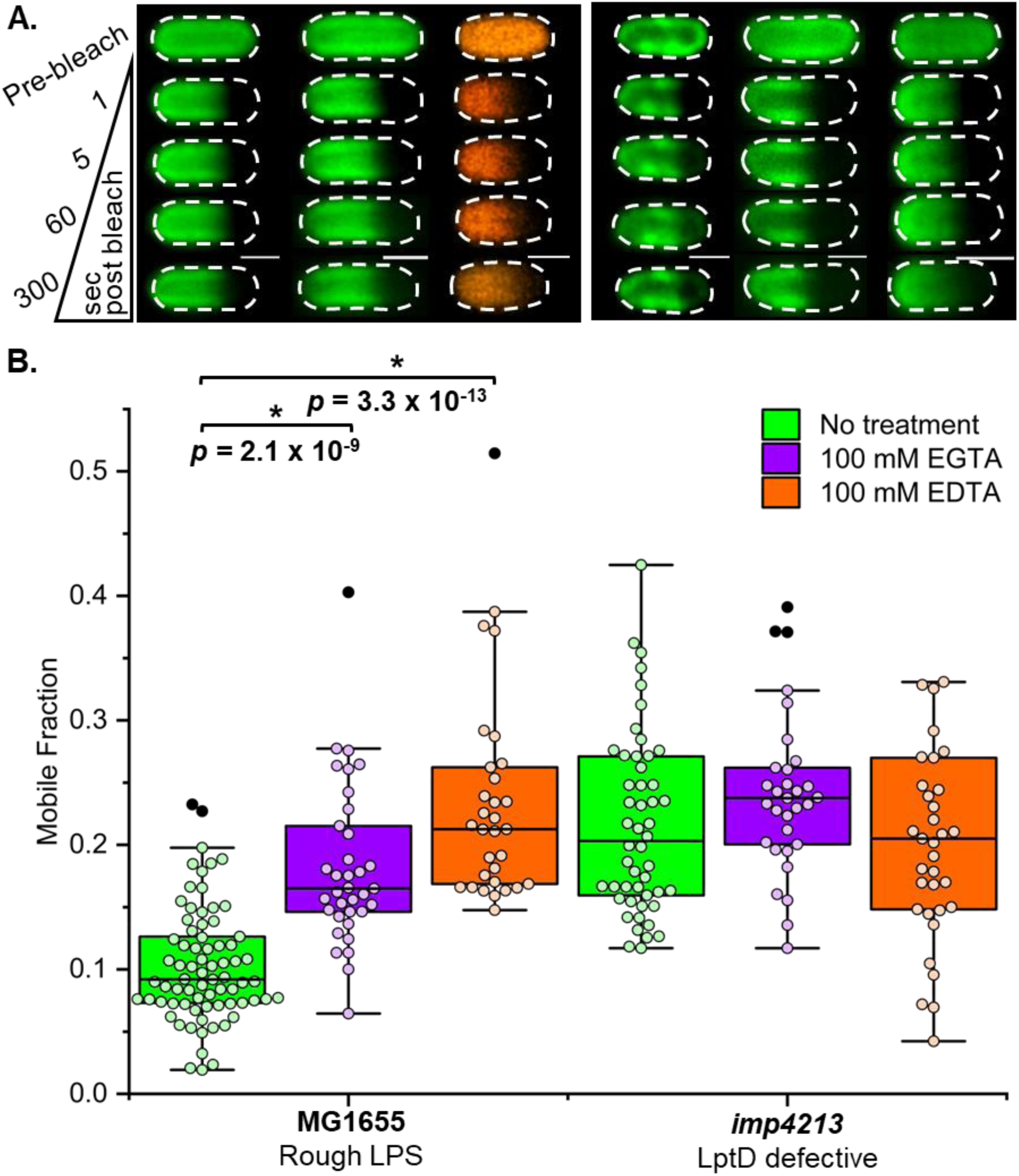
Observed differences in the sensitivity of LPS mobility to chelator treatments for *E. coli* MG1655 and *imp4213* cells suggests divalent cation-mediated LPS-LPS interactions contribute to LPS restriction in the OM. **A.** Representative FRAP sequences for AF488-LPS (green) or AF594-LPS (orange) in the OM of *E. coli* MG1655 and *imp4213* cells without (left-hand image sequence) or with 100 mM EGTA (middle image sequence) and 100 mM EDTA (right-hand image sequence) treatments. Scale bars: 1.0 µm. **B.** Effect of 100 mM EGTA and 100 mM EDTA treatments on LPS mobile fraction distributions in the OM of *E. coli* MG1655 and *imp4213* cells measured via FRAP. Boxes represent interquartile ranges with intersecting horizontal lines representing median values. MG1655: no treatment, median = 0.092 (n = 74); 100 mM EGTA, median = 0.165 (n = 35); 100 mM EDTA, median = 0.213 (n = 31). *imp4213*: no treatment, median = 0.203 (n = 50); 100 mM EGTA, median = 0.237 (n = 31); 100 mM EDTA, median = 0.205 (n = 35). The disruption of divalent-cation mediated LPS-LPS interactions by chelator treatments is responsible for the observed significant increase (by Mann-Whitney test) in LPS mobile fractions for the MG1655 cells. In contrast, the disruption of OM asymmetry in the *imp4213* cells caused by phospholipid flipping into the outer leaflet of the OM appears to disrupt these divalent-cation mediated interactions yielding a much lower overall sensitivity to chelator treatment for this strain.

**Supplementary Figure 8:**
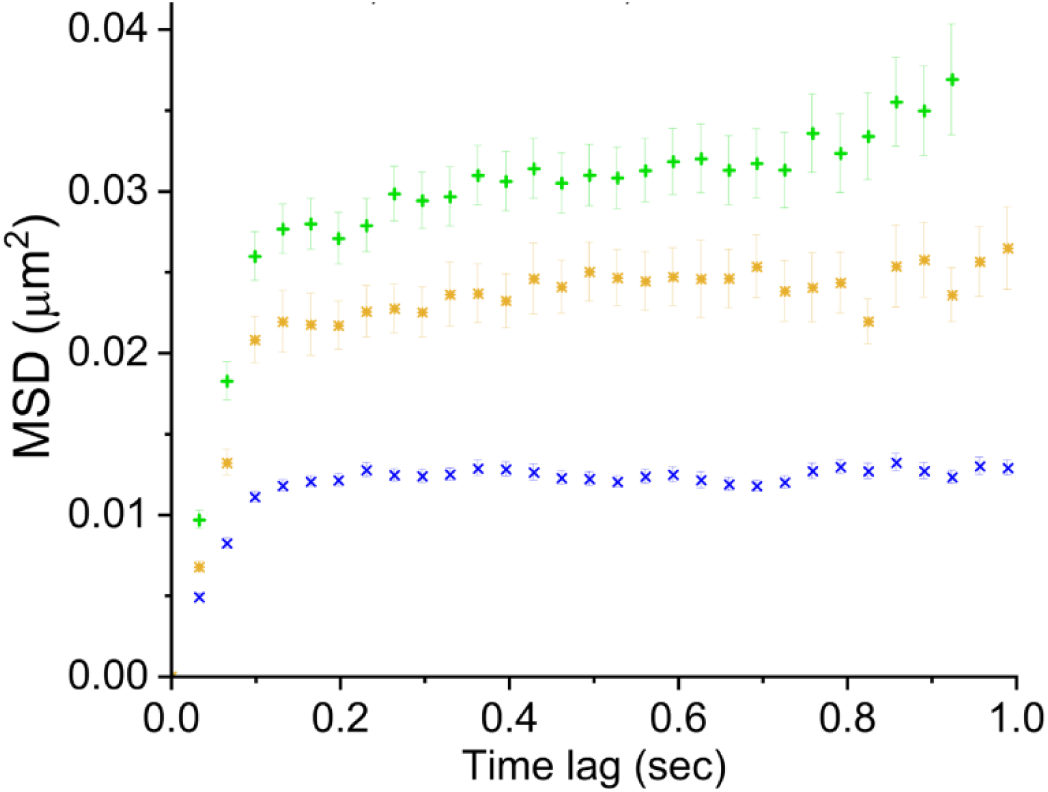
Effect of chelator treatments on the lateral diffusion of OMPs (CirA receptor) as measured on *E. coli* MG1655 cells by SPT-TIRFM. MSD was calculated for single AF488-labelled colicin Ia / CirA receptor complexes that could be tracked for at least 0.9 s before photobleaching (error is reported as S.E.M.) and were not immobilised on the quartz surface (MSD_end of trajectory_ > 0.008 µm^2^). All total internal reflection fluorescence microscopy video data was collected at 30 Hz from a minimum of three experimental replicates using 488 nm laser illumination. The MSD value for all OMP complexes on untreated *E. coli* MG1655 cells (blue **×**, n = 84) and on cells treated with 100 mM EDTA (green **+**, n = 99) or 100 mM EGTA (yellow ν, n = 74) approached an asymptotic value which was consistent with confined lateral diffusion. However, the level of confinement for OMP complexes was reduced in the chelator treated cells relative to the untreated cells. Linear regression of the MSD for the first 4 time delays yielded D_2D_ ≈ 0.0454 µm^2^/s and D_2D_ ≈ 0.0402 µm^2^/s, respectively, for the OMP complexes in EDTA- and EGTA-treated cells. These D_2D_ values were increased relative to the value for untreated cells (∼0.0178 µm^2^/s). The reduction in membrane confinement induced by chelator treatments was accompanied by a 2- to 3-fold increase in the apparent rate of lateral diffusion.

**Supplementary Figure 9:**
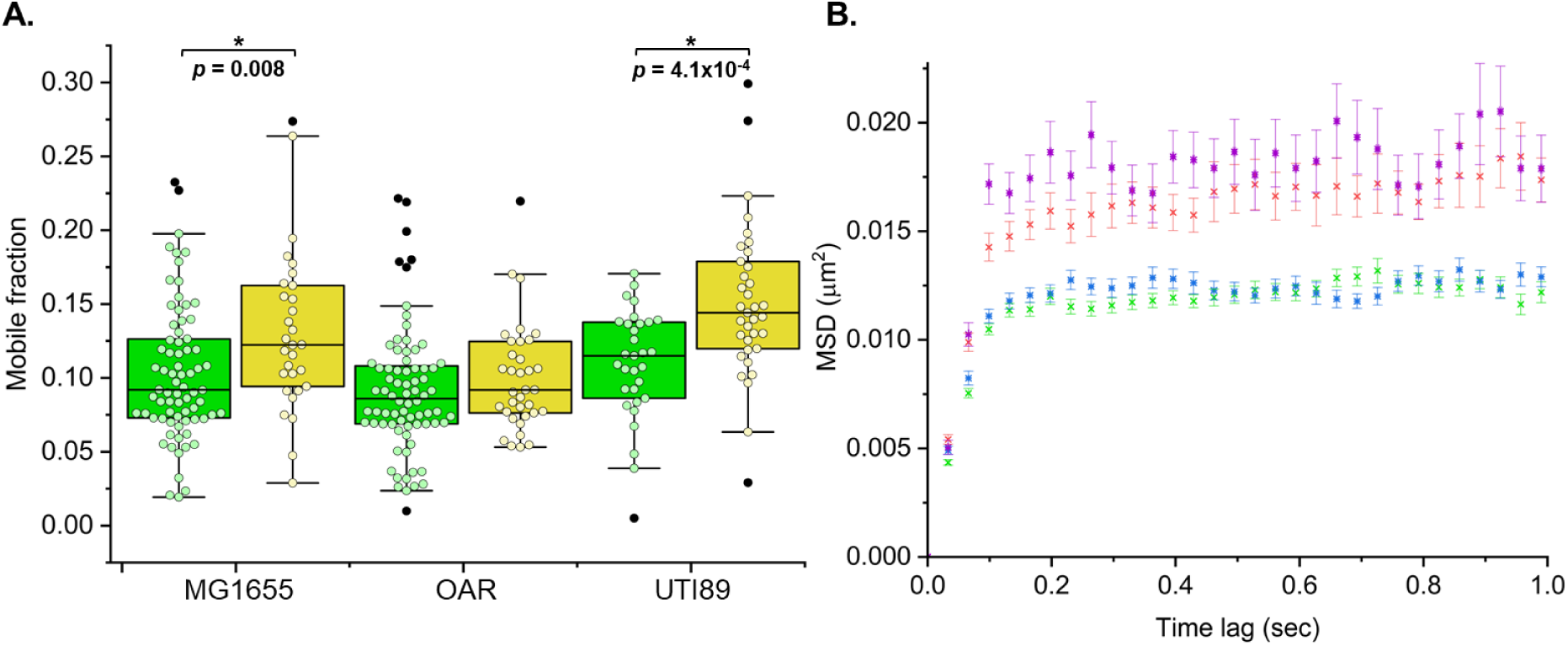
Treatment of bacterial cells with a non-lethal concentration of chaotropic agent (300 mM urea) induced a minor increase in lateral mobility of LPS and OMPs. **A.** Effect of 300 mM urea (a concentration comparable to that found in human urine) on LPS mobile fractions in the OM of *E. coli* MG1655, *E. coli* DFB1655 O-antigen restored (OAR) and uropathogenic *E. coli* UTI89 cells measured via FRAP. MG1655: no treatment (green box), median = 0.092 (n = 74); 300 mM urea treatment (yellow box), median = 0.122 (n = 30). OAR: no treatment (green box), median = 0.086 (n = 76); 300 mM urea treatment (yellow box), median = 0.092 (n = 35). UTI89: no treatment (green box), median = 0.115 (n = 31); 300 mM urea treatment (yellow box), median = 0.144 (n = 35). Significant increases in the LPS mobile fraction distribution were observed for the urea treatment of MG1655 cells (by Mann-Whitney test, *p* = 0.0082) and UTI89 cells (by Mann-Whitney test, *p* = 4.105 × 10^−4^). Whiskers represent maximum and minimum values, and each coloured dot represents an individual measurement collected over at least three experimental replicates. Dark grey dots outside whisker bounds are classified as outliers. **B.** Mean-squared displacement (MSD) was calculated for single AF488-labelled LPS molecules and single AF488-labelled colicin Ia / CirA receptor complexes that could be tracked for at least 1 s before photobleaching (error is reported as S.E.M.) and were not immobilised on the quartz surface (MSD_end of trajectory_ > 0.008 µm^2^). All total internal reflection fluorescence microscopy video data was collected at 30 Hz from a minimum of three experimental replicates using 488 nm laser illumination. The MSD for LPS molecules and the CirA receptor on untreated *E. coli* MG1655 cells and 300 mM urea treated cells approached an asymptotic value which was consistent with confined lateral diffusion. Treatment of cells with this chaotropic agent produced no measurable difference in the asymptotic MSD value relative to untreated cells for LPS and the CirA receptor. Linear regression of the MSD for the first 4 time delays yielded D_2D_ ≈ 0.0182 µm^2^/s (green **×**, n = 156) and D_2D_ ≈ 0.0246 µm^2^/s (red **×**, n = 154) for LPS on untreated and treated cells, respectively. Linear regression of the MSD for the same time delays yielded D_2D_ ≈ 0.0178 µm^2^/s (blue ®, n = 84) and D_2D_ ≈ 0.0319 µm^2^/s (purple ®, n = 107) for the CirA receptor on untreated and treated cells, respectively. The SPT data demonstrate that disruptions to the structure of the oligosaccharide domain induced by 300 mM urea cause a small reduction in the overall lateral confinement of LPS and CirA receptor, which is accompanied by a < 2-fold increase in the apparent rate of lateral diffusion for both LPS and the CirA receptor.

**Supplementary Figure 10:**
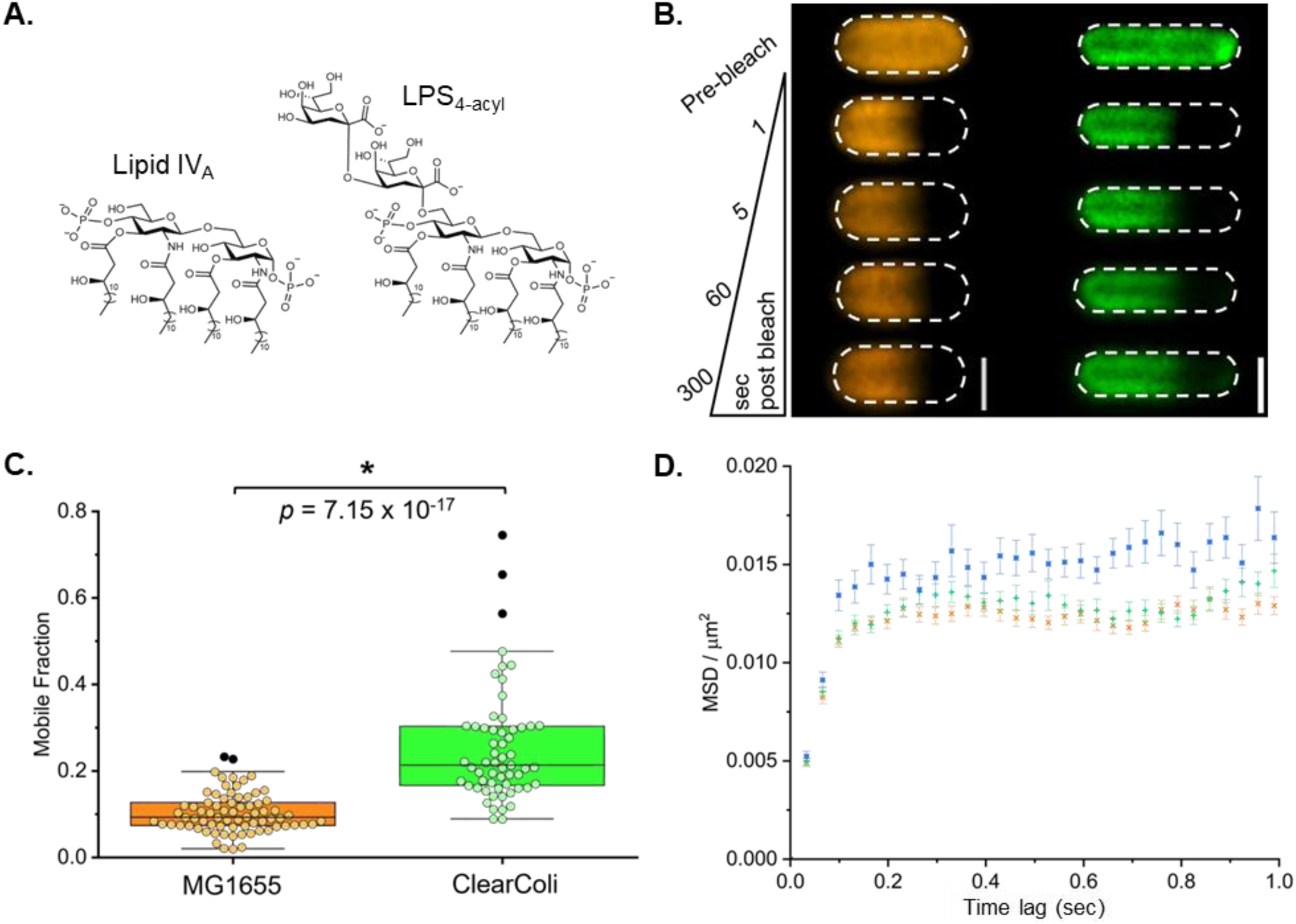
Reduction in acyl chain mediated intermolecular contacts between LPS molecules causes an increase in lateral mobility as detected by FRAP and SPT-TIRFM. **A.** ClearColi™ strain inserts an LPS-like molecule in the outer leaflet of the OM which is tetra-acylated Lipid IV_A_. Metabolic labelling with Kdo or Kdo-azide produces a deep rough-like LPS with 4 acyl chains (LPS_4-acyl_). The chemical structures of Lipid IV_A_ and deep rough-like LPS (LPS_4-acyl_) produced by ClearColi™ in the absence (Lipid IV_A_) or presence (LPS_4-acyl_) of metabolic labelling with Kdo sugar are presented here. **B.** Representative time-lapse FRAP images monitoring fluorescence recovery in the photobleached region of the OM for *E. coli* MG1655 cells with AF568-labelled rough LPS and ClearColi™ cells with AF488-labelled LPS_4-acyl_ (cells grown with 4 mM Kdo-azide). Scale bars: 1.0 µm. **C.** Lateral mobility of AF488- and AF568-labelled LPS measured by FRAP. LPS mobile fractions in the OM of *E. coli* MG1655 (rough LPS) and ClearColi™ grown with Kdo-azide (deep rough-like, tetra-acylated Lipid IV_A_). A statistically significant increase (by Mann-Whitney test, *p* = 7.147 × 10^−17^) in the LPS mobile fraction was observed in the ClearColi™ strain (median = 0.213, n = 59) relative to the wild-type *E. coli* MG1655 strain (median = 0.092, n = 74). **D.** Mean-squared displacement (MSD) was calculated for single AF488-labelled colicin Ia / CirA receptor complexes that could be tracked for at least 1 s before photobleaching (error is reported as S.E.M.) and were not immobilised on the quartz surface (MSD_end of trajectory_ > 0.008 µm^2^). All total internal reflection fluorescence microscopy video data was collected at 30 Hz from a minimum of three experimental replicates (except for ClearColi™ cells grown with 4 mM Kdo where n = 2) using 488 nm laser illumination. The MSD for all OMP complexes on *E. coli* MG1655 cells (yellow **×**, n = 84), ClearColi™ cells grown without Kdo sugar (green **+**, n = 90) and ClearColi™ cells grown with 4 mM Kdo sugar (blue **×**, n = 78) approached an asymptotic value, which was consistent with confined lateral diffusion. Linear regression of the MSD for the first 4 time delays yielded D_2D_ ≈ 0.0178 µm^2^/s, D_2D_ ≈ 0.0181 µm^2^/s and D_2D_ ≈ 0.0229 µm^2^/s, respectively, for these cells. These results demonstrate that changes in acyl chain mediated interactions between Lipid IV_A_ molecules in the outer leaflet of the OM primarily affect LPS-LPS interactions, with minimal effect on LPS-OMP interactions.

**Supplementary Figure 11.**
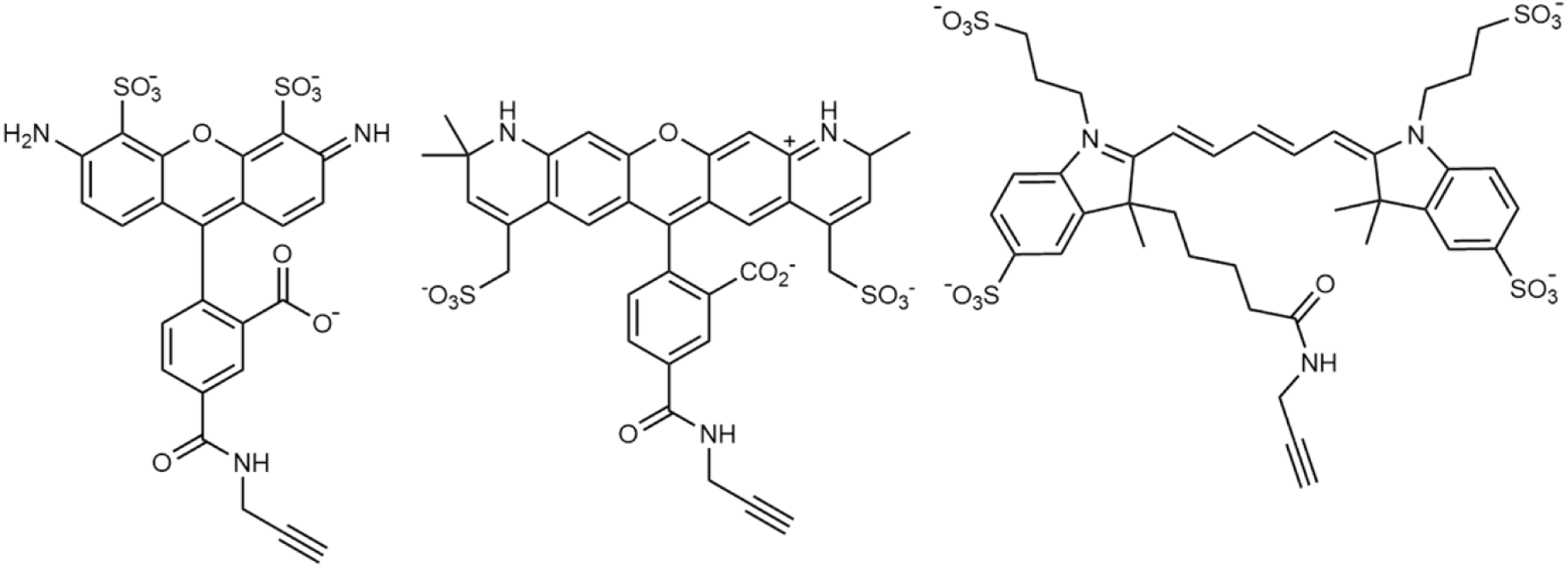
Structures of commercially available alkyne functionalised dyes: AF488-alkyne (left), AF568-alkyne (middle) and AZ647-alkyne (right).

**Supplementary Figure 12.**
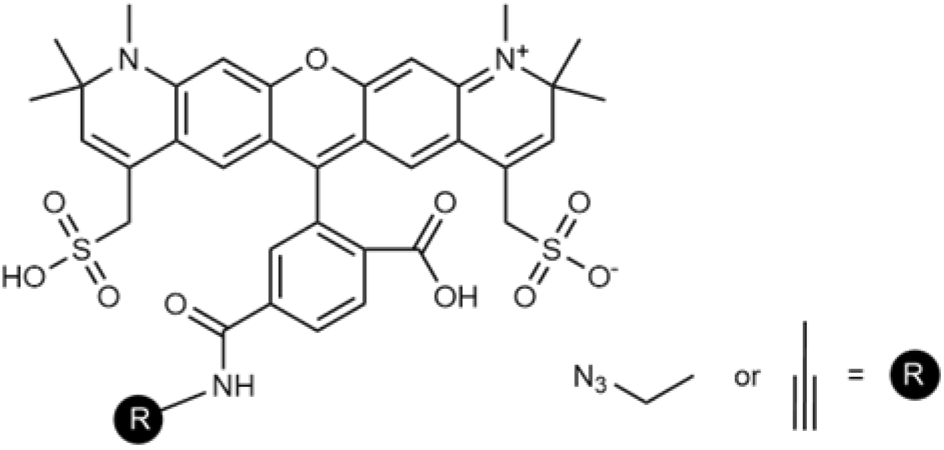
Structures of commercially available AZ594 azide / alkyne dyes.

**Supplementary Figure 13.**
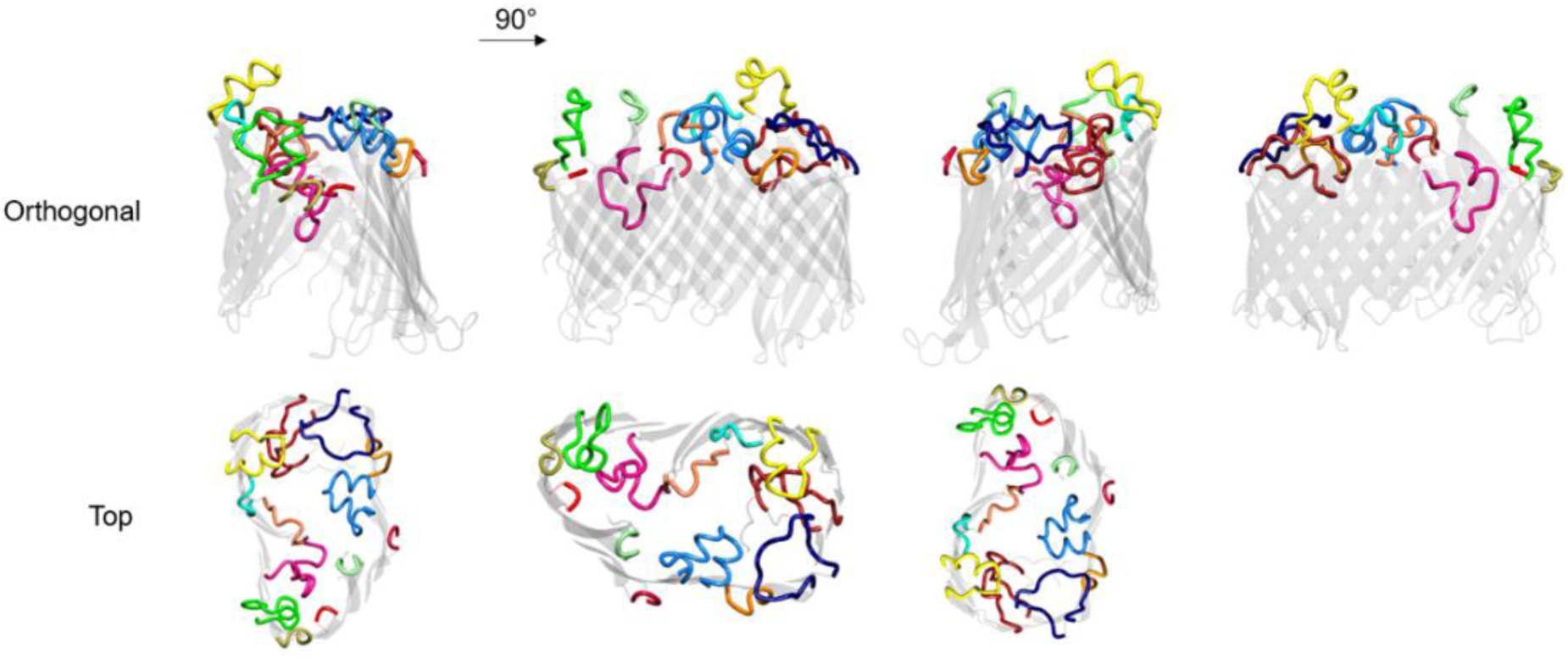
*Escherichia coli* K12 LptD crystal structure [PDB: 4RHB] highlighting extracellular loops investigated for possible ncAA incorporation via genetic code expansion (GCE).

**Supplementary Figure 14.**
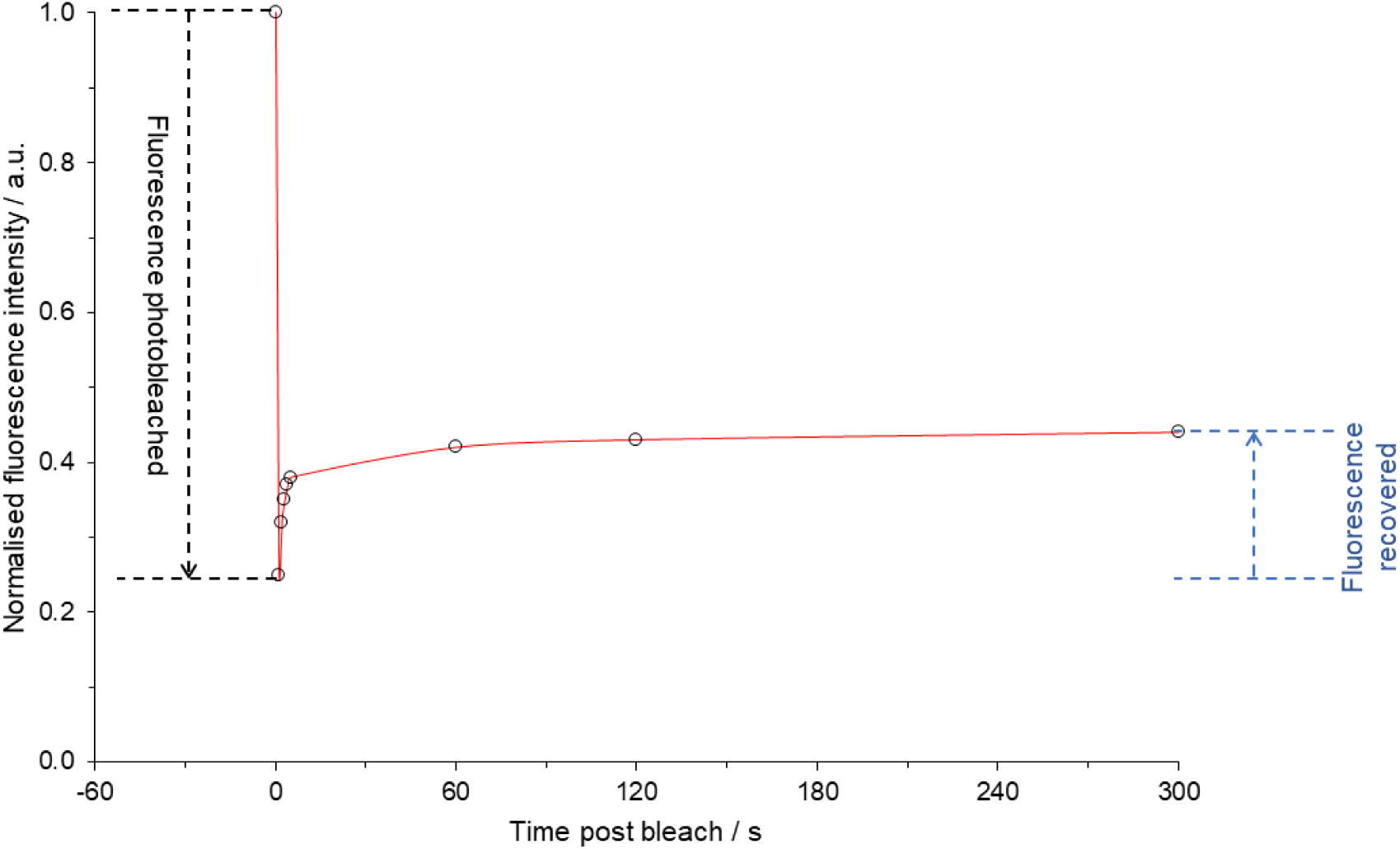
Example double-normalised FRAP recovery curve.

**Supplementary Table 1:**
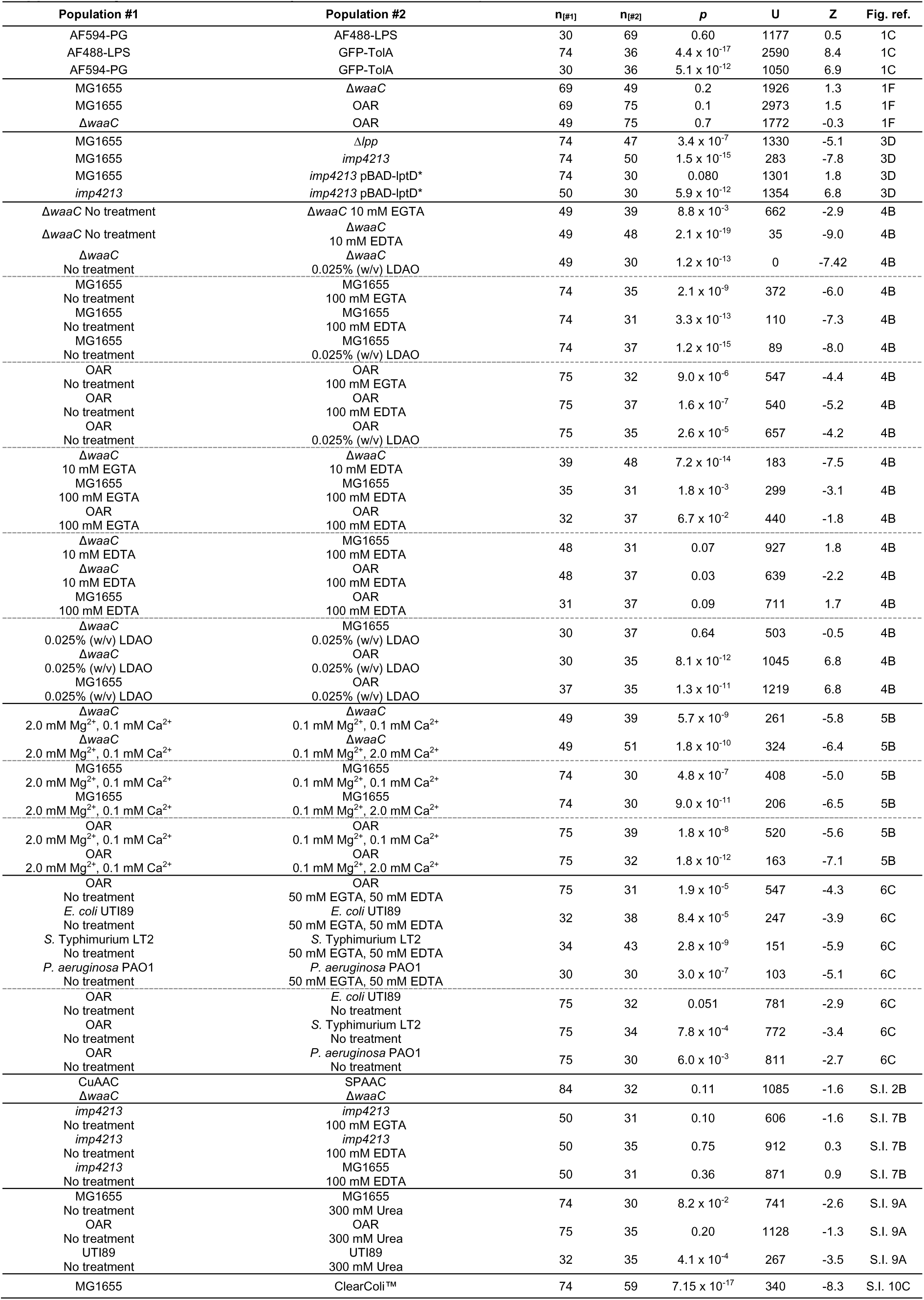
Mann-Whitney t-test statistics from comparison of FRAP-derived mobile fraction distributions.

**Supplementary Table 2:** Lateral diffusion of LPS and CirA receptor in the Gram-negative bacterial outer membrane characterised by SPT-TIRFM.

| Bacterial strain | Treatment | SPT probe | SPT imaging rate (Hz) | Number of tracks | D<sub>2D</sub> ± S.E.M. (μm<sup>2</sup>/s) | Confinement diameter ± S.E.M. (μm) | Type of lateral diffusion |
| --- | --- | --- | --- | --- | --- | --- | --- |
| *E. coli* MG1655 | None | AF488-rough LPS | 30 | 156 | 0.0182 ± 0.000645 | 0.543 ± 0.00666 | confined |
| *E. coli* MG1655 | None | AF488-rough LPS | 67 | 85 | 0.0621 ± 0.00399 | 0.662 ± 0.0139 | confined |
| *E. coli* MG1655 | 100 mM EDTA | AF488-rough LPS | 30 | 118 | 0.0377 ± 0.00266 | 0.815 ± 0.0238 | confined |
|  |  |  | 30 | 31 | 0.153 ± 0.0216 | --- | free |
| *E. coli* MG1655 | 100 mM EGTA | AF488-rough LPS | 30 | 108 | 0.0373 ± 0.00297 | 0.748 ± 0.0200 | confined |
|  |  |  | 30 | 21 | 0.113 ± 0.0143 | --- | free |
| *E. coli* MG1655 | 300 mM urea | AF488-rough LPS | 30 | 154 | 0.0246 ± 0.00136 | 0.613 ± 0.0139 | confined |
| *E. coli* MG1655 | 300 mM urea | AF488-rough LPS | 67 | 107 | 0.101 ± 0.00884 | 0.880 ± 0.0389 | confined |
| *E. coli* imp4213 | None | AF488-rough LPS | 30 | 104 | 0.0347 ± 0.00237 | 0.723 ± 0.0207 | confined |
|  |  |  | 30 | 49 | 0.0550 ± 0.00744 | --- | free |
| *E. coli* imp4213 | None | AF488-rough LPS | 67 | 78 | 0.0865 ± 0.00672 | 0.774 ± 0.0272 | confined |
|  |  |  | 67 | 14 | 0.140 ± 0.0254 | --- | free |
| *E. coli* ΔwaaC | None | AF488-deep rough LPS | 30 | 158 | 0.0182 ± 0.000722 | 0.542 ± 0.00617 | confined |
| *E. coli* ΔwaaC | None | AF488-deep rough LPS | 67 | 144 | 0.0480 ± 0.00266 | 0.596 ± 0.00981 | confined |
| *E. coli* MG1655 | None | AF488-colicin Ia / CirA | 30 | 84 | 0.0178 ± 0.000623 | 0.540 ± 0.00665 | confined |
| *E. coli* MG1655 | None | AF488-colicin Ia / CirA | 67 | 105 | 0.0514 ± 0.00242 | 0.583 ± 0.00837 | confined |
| *E. coli* MG1655 | 100 mM EDTA | AF488-colicin Ia / CirA | 30 | 99 | 0.0454 ± 0.00340 | 0.846 ± 0.0229 | confined |
| *E. coli* MG1655 | 100 mM EDTA | AF488-colicin Ia / CirA | 67 | 40 | 0.140 ± 0.0155 | 1.10 ± 0.0624 | confined |
| *E. coli* MG1655 | 100 mM EGTA | AF488-colicin Ia / CirA | 30 | 74 | 0.0402 ± 0.00471 | 0.740 ± 0.0216 | confined |
| *E. coli* MG1655 | 100 mM EGTA | AF488-colicin Ia / CirA | 67 | 92 | 0.0620 ± 0.00406 | 0.704 ± 0.0238 | confined |
| *E. coli* MG1655 | 300 mM urea | AF488-colicin Ia / CirA | 30 | 107 | 0.0319 ± 0.00222 | 0.636 ± 0.0181 | confined |
| *E. coli* MG1655 | 300 mM urea | AF488-colicin Ia / CirA | 67 | 104 | 0.0672 ± 0.00515 | 0.667 ± 0.0201 | confined |
| *E. coli* ClearColi | Cultured with 4 mM Kdo sugar | AF488-colicin Ia / CirA | 30 | 78 | 0.0229 ± 0.00206 | 0.600 ± 0.0124 | confined |
| *E. coli* ClearColi | Cultured without Kdo sugar | AF488-colicin Ia / CirA | 30 | 90 | 0.0181 ± 0.000820 | 0.546 ± 0.00760 | confined |

**Supplementary Table 3:** Cell numbers (n) for FRAP data and median mobile fraction values for cells cultured in M9 CDM with modified divalent cation concentrations (data plotted in Figure 5)

| Strain | M9 CDM type | Treatment | n | Median mobile fraction |
| --- | --- | --- | --- | --- |
| $\Delta waaC$ | high $Ca^{2+}$,<br>low $Mg^{2+}$ | none | 51 | 0.15 |
| $\Delta waaC$ | high $Ca^{2+}$,<br>low $Mg^{2+}$ | 5 mM EDTA, 5 mM EGTA | 39 | 0.21 |
| $\Delta waaC$ | low $Ca^{2+}$,<br>low $Mg^{2+}$ | none | 44 | 0.15 |
| $\Delta waaC$ | low $Ca^{2+}$,<br>low $Mg^{2+}$ | 5 mM EDTA, 5 mM EGTA | 35 | 0.19 |
| $\Delta waaC$ | low $Ca^{2+}$,<br>high $Mg^{2+}$ | none | 49 | 0.09 |
| $\Delta waaC$ | low $Ca^{2+}$,<br>high $Mg^{2+}$ | 5 mM EDTA, 5 mM EGTA | 30 | 0.24 |
| MG1655 | high $Ca^{2+}$,<br>low $Mg^{2+}$ | none | 30 | 0.18 |
| MG1655 | high $Ca^{2+}$,<br>low $Mg^{2+}$ | 50 mM EDTA, 50 mM EGTA | 58 | 0.23 |
| MG1655 | low $Ca^{2+}$,<br>low $Mg^{2+}$ | none | 34 | 0.18 |
| MG1655 | low $Ca^{2+}$,<br>low $Mg^{2+}$ | 50 mM EDTA, 50 mM EGTA | 31 | 0.17 |
| MG1655 | low $Ca^{2+}$,<br>high $Mg^{2+}$ | none | 74 | 0.09 |
| MG1655 | low $Ca^{2+}$,<br>high $Mg^{2+}$ | 50 mM EDTA, 50 mM EGTA | 31 | 0.18 |
| OAR | high $Ca^{2+}$,<br>low $Mg^{2+}$ | none | 32 | 0.20 |
| OAR | high $Ca^{2+}$,<br>low $Mg^{2+}$ | 50 mM EDTA, 50 mM EGTA | 42 | 0.20 |
| OAR | low $Ca^{2+}$,<br>low $Mg^{2+}$ | none | 39 | 0.15 |
| OAR | low $Ca^{2+}$,<br>low $Mg^{2+}$ | 50 mM EDTA, 50 mM EGTA | 30 | 0.16 |
| OAR | low $Ca^{2+}$,<br>high $Mg^{2+}$ | none | 75 | 0.08 |
| OAR | low $Ca^{2+}$,<br>high $Mg^{2+}$ | 50 mM EDTA, 50 mM EGTA | 31 | 0.17 |

**Supplementary Table 4:**
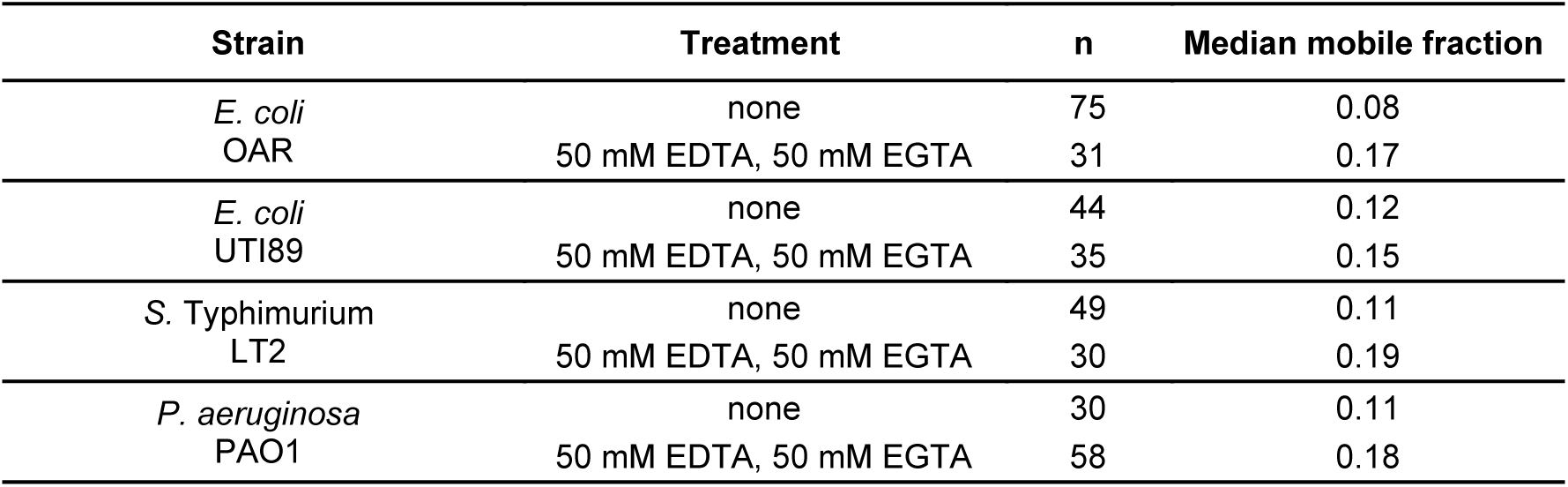
Cell numbers (n) for FRAP data and median mobile fraction values for bacterial cells with and without combined EDTA and EGTA treatment (data plotted in Figure 6)

**Supplementary Table 5.** Details of Gram-negative bacterial strains used in this study.

| Strain | Organism | LPS oligosaccharide length<sup>†</sup> | Selection antibiotic | Notes |
| --- | --- | --- | --- | --- |
| **K-12 substr. MG1655** | *E. coli* | Rough | None | Contains *rfb-50* mutation that results in absence of O-antigen synthesis. |
| **Δ*waaC*** | *E. coli* | Deep rough | 30 μg mL<sup>-1</sup> KAN | Keio collection strain JW3596-1. Δ*waaC* 733::Kan prohibits extension of LPS oligosaccharide domain beyond Kdo residues. |
| **ClearColi™** | *E. coli* | Endotoxin-free | None | Endotoxin-free strain (MG1655 background) with Lipid IV<sub>A</sub> as the only LPS-related glycolipid in the OM. This strain was only grown in LB-Miller medium. |
| **BW25113 GFP-TolA** | *E. coli* | Rough | 30 μg mL<sup>-1</sup> KAN | BW25113 strain transformed with pNP4 plasmid to produce GFP-tagged TolA as described previously. |
| **Δ*lpp*** | *E. coli* | Rough | 30 μg mL<sup>-1</sup> KAN | Keio collection strain JW1667-KC. |
| ***imp4213*** | *E. coli* | Rough | None | BW25113 background with *lptD* in-frame deletion (D330 – D352) mutation resulting in defective LptD function. |
| ***imp4213* pBAD-lptD\*** | *E. coli* | Rough | 100 μg mL<sup>-1</sup> AMP<br>35 μg mL<sup>-1</sup> CHL | *imp4213* transformed with pBAD-lptD* pEVOL-pylRS plasmids. Produces recombinant LptD with ncAA Lys analogue at position D600 when provided with the ncAA. |
| **Δ*ompA* pBAD-ompA\*** | *E. coli* | Rough | 100 μg mL<sup>-1</sup> AMP<br>35 μg mL<sup>-1</sup> CHL | Keio collection strain JW0940-KC transformed with pBAD-ompA* pEVOL-pylRS plasmid. Produces recombinant OmpA with ncAA Lys analogue at position E89 when provided with the ncAA. |
| **DFB1655 OAR** | *E. coli* | Smooth | 30 μg mL<sup>-1</sup> KAN | K12 substr. MG1655 with *wbbL* integrated downstream of IS5 element. Presents O16 serotype O-antigen. |
| **LT2** | *Salmonella enterica enterica* serovar Typhimurium strain LT2 | Smooth | None | Pathogenic model strain, serotype I 4,5,12:i:1,2. |
| **PAO1** | *Pseudomonas aeruginosa* | Smooth | None | Opportunistic pathogen model bacterial strain isolated from a wound in a patient. Capsule-forming serotype. |
| **UTI89** | *E. coli* | Smooth | None | Uropathogenic *E. coli* strain (UPEC) O18:K1:H7. Capsule-forming serotype. |

<sup>†</sup> LPS oligosaccharide type classifications: **Deep rough** = Lipid A-(Kdo)<sub>2</sub>, **Rough** = Lipid A-(Kdo)<sub>2</sub> plus core oligosaccharides, **Smooth** = Lipid A-(Kdo)<sub>2</sub> plus core oligosaccharides and 1 – 40 penta-saccharide O-antigen repeat units. **Antibiotic abbreviations:** **KAN** = Kanamycin (Sigma-Aldrich), **AMP** = Ampicillin (Melford Laboratories Ltd.), **CHL** = Chloramphenicol (Sigma-Aldrich)

**Supplementary Table 6.**
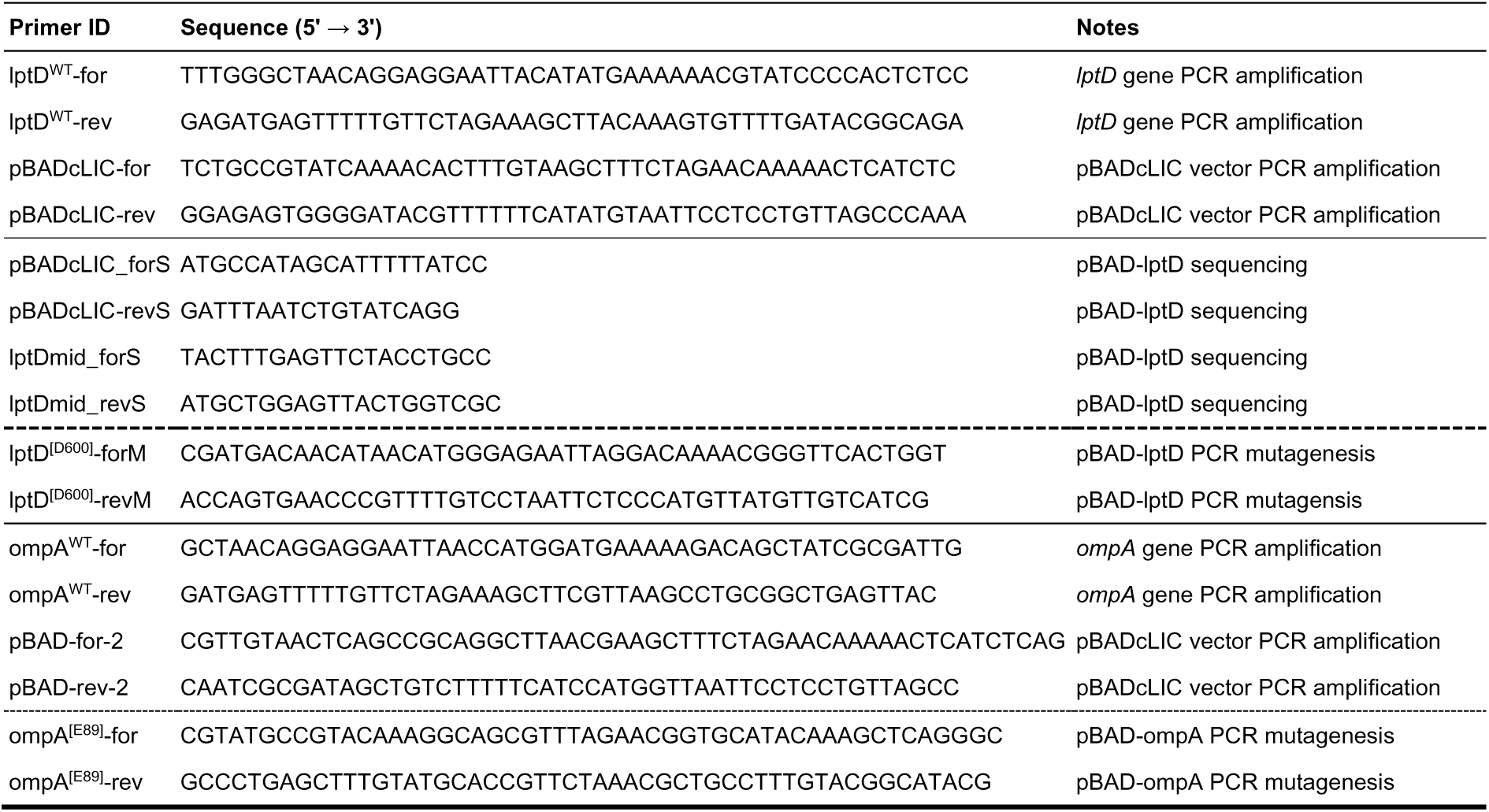
DNA primer sequences.

**Supplementary Table 7.** Composition of supplemented M9 chemically defined medium (pH 7.2)

| Component | Concentration |
| --- | --- |
| Na<sub>2</sub>HPO<sub>4</sub> | 48.0 mM |
| KH<sub>2</sub>PO<sub>4</sub> | 22.0 mM |
| NaCl | 08.6 mM |
| D-glucose | 0.4% (w/v) |
| NH<sub>4</sub>Cl | 1.0 g L<sup>-1</sup> |
| Casamino acids | 0.05% (w/v) |
| FeSO<sub>4</sub> | 0.1 mM |
| MgSO<sub>4</sub> | 2.0 mM |
| CaCl<sub>2</sub> | 0.1 mM |

**Supplementary Table 8:** Final concentrations of alkyne functionalised dyes (Click Chemistry Tools, Vector Laboratories, Inc.) used in CuAAC ‘Click-iT’ reaction mixtures.

| AZDye type | Final concentration (μM) |
| --- | --- |
| AF488 | 10 |
| AF568 | 20 |
| AZ647 | 50 |

**Supplementary Table 9.** Final concentrations of AZDye-DBCO dyes (Click Chemistry Tools, Vector Laboratories, Inc.) used in SPAAC reaction mixtures.

| AZDye DBCO type | Concentration (μM) |
| --- | --- |
| AZDye 488 | 25 |
| AZDye 568 | 50 |

**Supplementary Table 10.** Propargyl-OmpA^E89^ / Kdo-N_3_-LPS dual labelling combinations used to ensure fluorescent dye type and labelling order did not influence two-colour dSTORM results.

| Experimental replicate | First labelled species / fluorescent dye | Second labelled species / fluorescent dye |
| --- | --- | --- |
| 1 | LPS / AZ488-alkyne | OmpA<sup>E89</sup> / AZ647-N<sub>3</sub> |
| 2 | OmpA<sup>E89</sup> / AZ647-N<sub>3</sub> | LPS / AZ488-alkyne |
| 3 | LPS / AZ647-alkyne | OmpA<sup>E89</sup> / AZ488-N<sub>3</sub> |
| 4 | OmpA<sup>E89</sup> / AZ488-N<sub>3</sub> | LPS / AZ647-alkyne |

**Supplementary Table 11.** 4x TSDS-PAGE sample loading buffer composition.

| Component | Final concentration |
| --- | --- |
| Tris-Cl pH 6.8 | 0.25 M |
| Sodium dodecyl sulphate (SDS) | 227 mM |
| Bromophenol blue | 0.02 % (w/v) |
| Glycerol | 4.3 M |

**Note:** β-ME added to a final concentration 2% (v/v) immediately prior to its usage.

**Supplementary Table 12.** Concentrations of chelator, detergent and urea used for pre-FRAP treatments.

| LPS oligosaccharide glycoform | EGTA conc. (mM) | EDTA conc. (mM) | EGTA / EDTA conc. (mM) | LDAO conc. (% w/v) | Urea conc. (mM) |
| --- | --- | --- | --- | --- | --- |
| Deep rough | 10 | 10 | 5 / 5 | 0.025 | 300 |
| Rough | 100 | 100 | 50 / 50 | 0.025 | 300 |
| Smooth | 100 | 100 | 50 / 50 | 0.025 | 300 |

**Supplementary Table 13.** Composition of supplemented M9 chemically defined medium (CDM), supplemented M9 CDM with reduced [Mg^2+^] and supplemented M9 CDM with elevated [Ca^2+^] (differences across the three media types are highlighted in bold).

| Component | Standard M9 CDM<br>pH 7.2 (high [Mg<sup>2+</sup>]) | Low [Mg<sup>2+</sup>] M9 CDM<br>pH 7.2 | High [Ca<sup>2+</sup>] M9 CDM<br>pH 7.2 |
| --- | --- | --- | --- |
| NaCl | 0.5 g L<sup>-1</sup> | 0.5 g L<sup>-1</sup> | 0.5 g L<sup>-1</sup> |
| Na<sub>2</sub>HPO<sub>4</sub> | 6.78 g L<sup>-1</sup> | 6.78 g L<sup>-1</sup> | 6.78 g L<sup>-1</sup> |
| KH<sub>2</sub>PO<sub>4</sub> | 3.0 g L<sup>-1</sup> | 3.0 g L<sup>-1</sup> | 3.0 g L<sup>-1</sup> |
| NH<sub>4</sub>Cl | 1.0 g L<sup>-1</sup> | 1.0 g L<sup>-1</sup> | 1.0 g L<sup>-1</sup> |
| casamino acids | 0.05% [w/v] | 0.05% [w/v] | 0.05% [w/v] |
| D-glucose | 22 mM | 22 mM | 22 mM |
| **MgSO<sub>4</sub>** | 2 mM | **100 μM** | 100 μM |
| **CaCl<sub>2</sub>** | 100 μM | 100 μM | **2 mM** |
| FeSO<sub>4</sub> | 100 μM | 100 μM | 100 μM |

**Supplementary Table 14.**
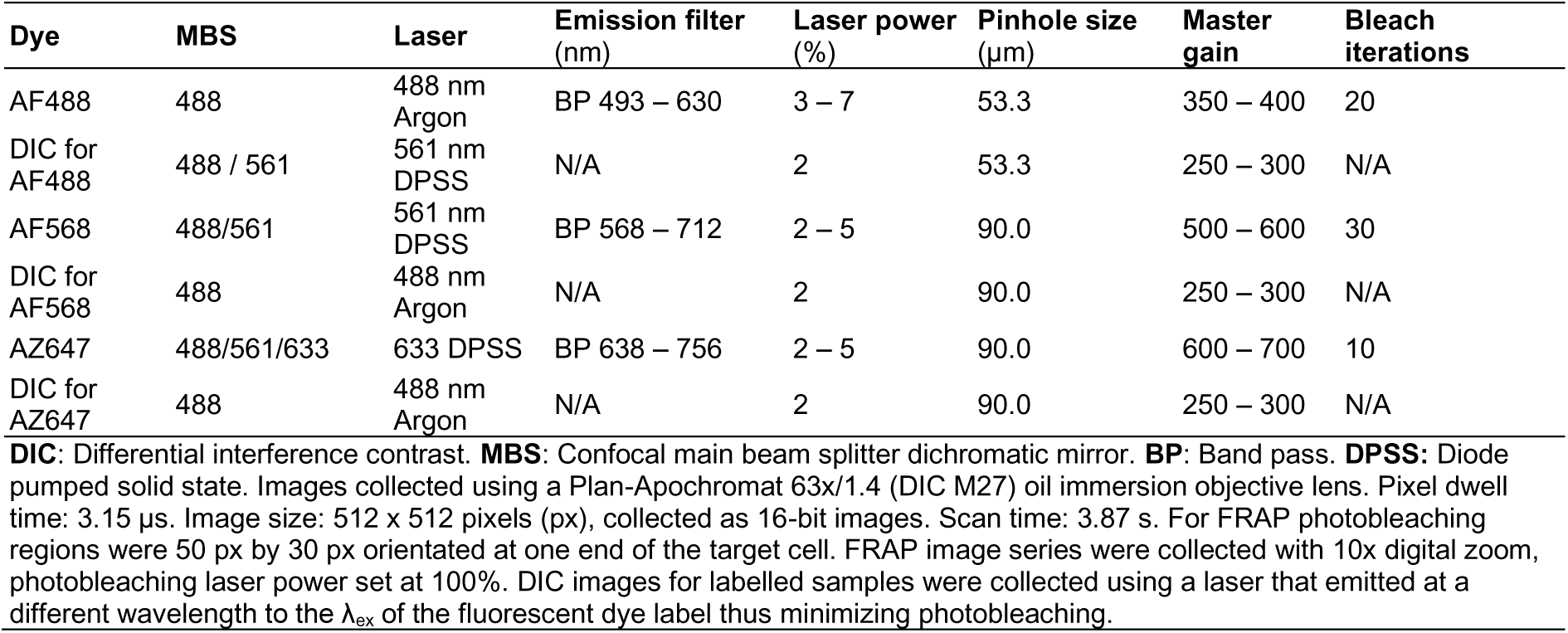
Upright Zeiss LSM710 confocal microscope settings.

**Supplementary Table 15.** Inverted Zeiss LSM780 confocal multiphoton microscope settings.

| Dye | MBS | Laser | Emission filter (nm) | Laser power (%) | Pinhole size (μm) | Master gain | Bleach iterations |
| --- | --- | --- | --- | --- | --- | --- | --- |
| AF488 | 488 / 561 | 488 nm Argon | BP 493 – 630 | 2 – 3 | 47.7 | 700 – 750 | 20 |
| DIC for AF488 | 488 / 561 | 561 nm DPSS | N/A | 2 | 53.3 | 250 – 300 | N/A |
| AF568 | 488 / 561 | 561 nm DPSS | BP 568 – 712 | 2 – 5 | 90.0 | 400 – 500 | 30 |
| DIC for AF568 | 488 | 488 nm Argon | N/A | 2 | 90.0 | 250 – 300 | N/A |

**DIC:** Differential interference contrast. **MBS:** Confocal main beam splitter dichromatic mirror. **BP:** Band pass. **HeNe:** Helium-neon. Images collected using a Plan-Apochromat 63x/1.4 (DIC M27) oil immersion objective lens. Pixel dwell time: 1.58 μs. Image size: 512 x 512 pixels, collected as 16-bit images. Scan time: 1.94 s.

**Supplementary Table 16.** Components in Glucose oxidase (GluOx) / Catalase (Cat) oxygen scavenging buffer system composition.

| Component | Volume (μL) |
| --- | --- |
| Degassed deionised H<sub>2</sub>O | 895 |
| PBS pH 7.4 [40 x] (BioStatus) | 25 |
| GluOx [10 mg mL<sup>-1</sup>] | 20 |
| Cat [2 mg mL<sup>-1</sup>] | 20 |
| Glucose [300 mg mL<sup>-1</sup>] | 40 |

**Supplementary Table 17.** Additional settings for Zeiss Elyra 7 SRM microscope 2D dSTORM image Acquisition.

| Dye | Laser beam splitter | Laser | Emission filter (nm) | xy scaling ($\mu\text{m pixel}^{-1}$) | TIRF mirror angle | TIRF collimator |
| --- | --- | --- | --- | --- | --- | --- |
| AZ488 | 405/488/561/642 | 488 nm DPSS | BP420-480 + BP490-550 | 0.097 | 60° | 230 |
| AZ647 | 405/488/561/642 | 561 nm DPSS | BP570-620 + LP655 | 0.097 | 60° | 400 |

**Supplementary Table 18.** Typical dSTORM image filtering settings.

| Filter | AF488 dye | AZ647 dye |
| --- | --- | --- |
| Precision (nm) | 1 – 60 | 1 – 60 |
| Number of photons | 1 – 2000 | 1 – 1250 |
| Point squared function half width (nm) | 60 – 200 | 100 – 300 |
| Background variance | 1 – 300 | 1 – 300 |
| Chi-squared | 0.4 – 1.2 | 0.4 – 1.2 |

